# Fibrotic lung-derived sphingosine-1-phosphate drives emotion-like disorders by regulating hippocampal neuroinflammation and cell death

**DOI:** 10.64898/2026.04.28.721267

**Authors:** Zhihao Xu, Deshang Xu, Keqin Liu, Yijia Yang, Xingjie Han, Fen Yang, Wenbin Nan, Jiang Du, Rui Guo, Yonghai Li, Ping Zhang, Yanli Liu, Juntang Lin

**Author notes:** Corresponding author: Zhihao Xu, PhD, Yanli Liu, PhD, Juntang Lin, PhD. These authors contributed equally to this work (Zhihao Xu, Deshang Xu, Keqin Liu).

## Abstract

The incidence of emotional disorders in patients with idiopathic pulmonary fibrosis (IPF) is substantially higher than that in the general population, severely compromising their quality of life. However, the underlying mechanisms remain poorly understood. In this study with multi-omics, we demonstrated that sphingosine-1-phosphate (S1P) derived from IPF lungs drive anxiety and depressive-like behaviors. Mechanistically, circulating S1P in the blood bound to hippocampal S1PR1 to regulate the PI3K/PKA/CREB signaling pathway, leading to synapse damage, the activation of microglia and astrocytes, neuroinflammation and ferroptosis in the hippocampus. Pharmacological inhibition of Sphk1, a key enzyme in S1P synthesis, reduced serum S1P levels and alleviated IPF-induced anxiety and depressive-like behaviors. Similarly, selective inhibition of hippocampal S1P receptor signaling using Fingolimod also attenuated neuroinflammation and ferroptosis and ameliorated mood disorders in IPF models. Collectively, these findings demonstrate that metabolite S1P from fibrotic lungs serves as a mediator of lung-to-brain functional influence, providing new insights into the IPF comorbid mood disorders and potential therapeutic targets.

## Introduction

Idiopathic pulmonary fibrosis (IPF) is a disease characterized by pulmonary interstitial fibrosis and progressive decline in lung function that severely impairs patients’ quality of life, affecting approximately 3 million people worldwide [1, 2]. Accumulating evidence indicates that the incidence of emotional disorders such as depression and anxiety, which are often clinically overlooked, is significantly higher in IPF patients than in the general population [3–5]. The prevalence of depression among IPF patients increases by 3.24% each year after diagnosis [6], suggesting that both physiological changes and the psychological stress following an IPF diagnosis may trigger the onset of mood disorders. The emergence of mood symptoms in IPF patients can adversely affect treatment adherence, reduce the efficacy of IPF-targeted therapies, and even increase the risk of suicidal ideation [4, 6, 7]. Therefore, research and interventions targeting comorbid mood disorders are highly important for the management of IPF.

Multiple factors may contribute to the development of mood comorbidity in IPF, including psychological stress, hypoxia, neural transmission, and circulating factors [8, 9]. In patients who develop new-onset depression after an IPF diagnosis, the mortality risk is significantly increased, and the cumulative initiation rate of antifibrotic therapy is markedly reduced. In contrast, pre-existing depression before diagnosis is not significant associated with mortality risk or the treatment initiation rate [6], suggesting that intervention for depressive symptoms should be a priority in IPF management. The brain has the highest oxygen demand of any organ. Severe fibrosis impairs pulmonary gas exchange and leads to complications such as hypoxemia and hypercapnia, which may represent the most direct pathway through which advanced lung disease affects brain function. However, some patients develop mood disorders even in the absence of hypoxia, indicating that hypoxia is not the sole cause [10, 11]. Pulmonary irritation and lesions may act on the sympathetic and sensory nerves innervating the lungs, transmitting signals rapidly to the brain and brainstem via long-distance axonal transport. Repeated or chronic stimulation from the lungs can thereby induce mood symptoms through neural pathways [12–14]. Intervening in the nerves that innervate the lungs can relieve emotional tension and depressive symptoms [15, 16]. Circulating factors derived from lung tissue—including cytokines, proteins and exosomes—may also affect brain function. Lung infection or injury releases large amounts of inflammatory cytokines such as IL-1β, IL-6, and TNF-α, which enter the bloodstream, compromise the blood–brain barrier, and subsequently trigger or exacerbate neuroinflammation in the central nervous system [11, 17]. Exosomes derived from lung cells may further influence neuronal function by transferring small RNA molecules [18]. Although antifibrotic therapy can alleviate mood symptoms, no drug is currently available to specifically improve mood disorder comorbidity in IPF patients [19]. These findings indicate that additional factors drive mood disorders in IPF patients and highlight the need to develop treatments targeting mood disorders in this patient population.

In patients with pulmonary fibrosis, pulmonary metabolism undergoes profound changes, including disruptions in glucose, lipid, and amino acid metabolism, all of which collectively drive the progression of IPF [20–22]. Among these, lipid metabolism disorder is one of the most prominent features of pulmonary fibrosis, characterized by significantly elevated levels of non-esterified fatty acids, long-chain acylcarnitines, and specific ceramides. These metabolites may serve as biomarkers of disease progression and prognosis in pulmonary fibrosis and could also represent potential therapeutic targets [23, 24]. Pulmonary metabolites may affect brain function via the bloodstream, leading to blood–brain barrier disruption, neuroinflammation, and cognitive impairment [25–29]. However, whether IPF-derived pulmonary metabolites that induce mood disorders exist has not yet been reported and identified.

Based on the above findings, we hypothesized that IPF-derived metabolites may induce the onset of emotional disorders and sought to identify such metabolites. To this end, we established a mouse model of pulmonary fibrosis and confirmed the presence of emotion-like disorders. Using liquid chromatograph mass spectrometer (LC-MS), we measured metabolites in lung tissue and serum, and performed transcriptomic sequencing of both lung and hippocampus. We found that elevated serum S1P, which was derived from the fibrotic lung, drive mood symptoms in IPF mice. We observed synapse damage, inflammation and cell death, increased expression of S1PR1, and dysregulation of the PI3K/PKA/CREB signaling pathway in the hippocampus. Pharmacological inhibition of S1P synthesis in the lung, or functional blockade of S1PR1 and its downstream pathway in the hippocampus, alleviated mood disorders in IPF mice. Our study elucidates the driving role of S1P derived from fibrotic lungs in mood disorders and provides new insights into the mechanisms and integrated treatment of emotional disorders induced by pulmonary diseases.

## Materials and methods

### Drugs, antibodies and kits

Bleomycin (BLM, HY-108345), S1P (HY-108496), SKI-V (HY-12895), and Fingolimod (FTY720, HY-11063) were purchased from MedChemExpress (MCE, Shanghai, China). The primary antibodies were listed in Supplementary Material 2. The secondary antibodies conjugated to horseradish peroxidase (EK1002), Enhanced Chemiluminescence Method Kit (EK1002) and protein quantitative bicinchoninic acid (BCA) kit (AR1189) were purchased from Boster Biological Technology Co., Ltd (Wuhan, China). The S1P ELISA kit (U96-3489E) was purchased from YOBIBIO Biological Technology Co., Ltd. (Shanghai, China), and the ELISA kits for the cytokines IL-1β (EK0394), IL-6 (EK0411), TNF-α (EK0527), CCL2 (EK0568), IL-4 (EK0405), and IL-10 (EK0417) were purchased from BOSTER Biological Technology Co., Ltd. (Wuhan, China).

### Animal and drug administration

Male C57BL/6 mice (8 weeks old, body weight 20–25 g) were purchased from Beijing Vital River Laboratory Animal Technology Co., Ltd. (Beijing, China). Prior to model establishment or drug administration, the mice were housed under standard experimental conditions for 7 days: 22 ± 2°C temperature, 50–60% relative humidity, and a 12 h light/dark cycle (lights on at 8:00 a.m.), with free access to food and water. The IPF model was induced via a single intranasal instillation of 1.5 U/kg bleomycin dissolved in a total of 50 μL of saline. S1P, dissolved in DMSO and diluted with saline, was administered intraperitoneally at a dose of 5 mg/kg body weight daily. SKI-V, dissolved in DMSO and diluted with saline, was administered intraperitoneally at a dose of 50 mg/kg body weight every other day. Fingolimod, dissolved in DMSO and diluted with saline, was administered intraperitoneally at a dose of 2 mg/kg body weight daily. All experimental protocols and animal care and use were approved by the Ethics Committee of Henan Medical University.

### Behavioral tests

Before behavioral testing, the mice were allowed to acclimate to the testing room for 1 hour.

### Open field test (OFT)

The apparatus (Xinruan, Shanghai, China) consisted of a square open field made of opaque acrylic (50 cm × 50 cm × 40 cm), illuminated with low light (approximately 20 lux in the center). The floor was divided into nine equal grids. Each mouse was gently placed in the central area and allowed to explore freely for 5 minutes. Behavior was recorded by an overhead VisuTrack camera (Xinruan, Shanghai, China). The time spent in the central grid was used as an indicator of anxiety-like behavior. After each test, the apparatus was thoroughly cleaned with 70% ethanol and dried to eliminate olfactory cues.

### Sucrose preference test (SPT)

During the entire testing period, the mice were housed individually. After 4 hours of food and water deprivation, each mouse was simultaneously given two pre-weighed bottles (one containing 1% sucrose solution and the other containing tap water) for 24 hours. Liquid consumption was calculated by weighing the bottles before and after the test. The sucrose preference index was calculated as follow: sucrose preference (%) = [sucrose solution intake / (sucrose solution intake + water intake)] × 100%.

### Tail suspension test (TST)

Each mouse was fixed with adhesive tape approximately 1 cm from the tip of the tail and suspended upside down from a horizontal bar inside a sound-attenuating chamber (30 cm × 30 cm × 60 cm, three sides opaque). Each mouse’s head was positioned about 10 cm above the floor of the chamber. The test lasted for 6 minutes, and the immobility time during the last 4 minutes was analyzed. Immobility was defined as the absence of any active movement, with the mouse hanging passively and vertically. Behavior was recorded and analyzed by VisuTrack camera (Xinruan, Shanghai, China). The apparatus was cleaned with 70% ethanol after each test.

### Forced swimming test (FST)

The apparatus (Xinruan, Shanghai, China) consisted of a transparent organic glass cylinder (30 cm in height, 20 cm in diameter) filled with water (23–25°C) to a depth of 15 cm, ensuring that the mouse’s tail or hind limbs could not touch the bottom. Each mouse was gently placed into the water and tested for 6 minutes. Behavior was recorded and analyzed by a VisuTrack camera (Xinruan, Shanghai, China). The immobility time during the last 4 minutes was quantified. Immobility was defined as the mouse making only minimal movements necessary to keep its head above water without actively struggling or swimming. At the end of the test, the mouse was removed, gently dried with a soft towel, placed under a heat lamp for 15 minutes to recover, and then returned to its home cage. The water was changed after each test to eliminate olfactory cues and maintain a constant water temperature.

### Sampling

Following the behavioral tests, blood was collected from the retro-orbital sinus of each mouse. The blood samples were allowed to stand at 4°C for 24 hours, and then the serum was separated and centrifuged at 12,000 rpm for 15 minutes at 4°C. The serum was stored at −20°C for subsequent metabolite measurement and ELISA. The mice were euthanized by cervical dislocation. Lung tissues were harvested and rinsed to remove residual blood. Lung tissue was fixed in 4% paraformaldehyde for sectioning and subsequent hematoxylin and eosin (H&E), and Masson staining. The lungs were frozen for transcriptomic sequencing, metabolomic profiling, ELISA, qPCR, and Western blotting. The brains were collected and rinsed to remove residual blood. Whole brains were fixed in 4% paraformaldehyde for sectioning and immunofluorescence staining. For Golgi staining, the brains were immersed in Golgi fixative (G1069, Servicebio, Wuhan, China). Hippocampi were dissected, immediately frozen, and stored at −80°C for transcriptomic sequencing, ELISA, qPCR, and Western blotting.

### Histological staining and analysis

#### H&E staining

The lungs were fixed in 4% paraformaldehyde at 4°C for 48 hours, dehydrated through a graded ethanol series, cleared in xylene, and infiltrated with molten paraffin. The tissues were subsequently embedded in paraffin blocks and allowed to solidify at room temperature. The paraffin blocks were trimmed and sectioned continuously at a thickness of 3 μm using a microtome. The sections were flattened in a 40°C water bath, placed onto adhesive slides, and baked at 60°C for 2 hours. The sections were then deparaffinized, rehydrated through a graded ethanol series, and rinsed with distilled water for 2 minutes. The sections were immersed in Harris hematoxylin solution for 5–8 minutes, and excess stain was removed with distilled water. They were differentiated in 1% hydrochloric acid in ethanol for 2–5 seconds, immediately followed by rinsing under running tap water for 15 minutes. The sections were then immersed in 0.5% eosin aqueous solution for 1–3 minutes, quickly rinsed with distilled water, dehydrated in ethanol, cleared in xylene, mounted with neutral resin, and coverslipped. The slides were air-dried at room temperature and observed under a microscope for image acquisition.

#### Masson staining

Paraffin-embedded lung tissue sections were stained using a Masson’s trichrome staining kit (G1006, Servicebio, Wuhan, China) according to the manufacturer’s instructions. The detailed procedure was as follows. The sections were immersed in 2.5% potassium dichromate solution, placed in a 65°C oven for 30 minutes, and then rinsed under running tap water. The sections were stained with a 1:1 mixture of Wiegert’s hematoxylin solution for 1 minute, washed with tap water, differentiated with differentiation solution for a few seconds, and rinsed again with tap water. The sections were then immersed in preheated (55°C) Ponceau-acid fuchsin solution for 6 minutes, rinsed with tap water, treated with 1% phosphomolybdic acid for 1 minute, and immediately transferred to 2.5% aniline blue solution at 55°C for 2–30 seconds. The sections were rinsed and differentiated with 1% acetic acid, dehydrated in absolute ethanol, cleared in xylene, and mounted with neutral resin. Observations and image acquisition were performed under a microscope. The collagen appeared blue. Relative collagen content was quantified using ImageJ v1.54p.

#### Immunofluorescence Staining

The brains were fixed for 72 hours, and then dehydrated through a graded sucrose solution. The tissues were embedded in optimal cutting temperature (OCT) compound (G6059, Servicebio, Wuhan, China) and sectioned using a cryostat (Leica). Sections were cut at a thickness of 20 μm, mounted onto adhesive slides, and baked in a 37°C oven for 10 minutes to remove excess moisture. The sections were fixed in methanol for 30 minutes and permeabilized with 0.5% Triton X-100 (diluted in PBS) for 30 minutes. After washing, the sections were blocked with 5% BSA at room temperature for 1 hour. The sections were then incubated with primary antibodies in a humidified chamber at 4°C overnight, followed by incubation with the corresponding secondary antibodies at room temperature for 1 hour in the dark. DAPI solution was added and the sections were incubated at room temperature for 30 minutes in the dark. An autofluorescence quencher (G1221) was applied for 5 minutes, and the sections were mounted with anti-fade mounting medium. The sections were scanned using a digital slide scanner (3DHISTECH, Pannoramic MIDI). The fluorescence signal intensity and density were quantified using ImageJ v1.54p. Microglial and astrocytic skeleton morphology and branch complexity were analyzed using the skeletonize tool and Sholl analysis plugin of ImageJ v1.54p, respectively.

#### Golgi Staining

Mouse brain tissues were fixed in Golgi fixative for 3 days, then completely immersed in Golgi staining solution and placed in a cool, ventilated, dark environment for 14 days. The staining solution was replaced every 3 days. The tissues were sectioned at 60 μm thickness using a vibratome, mounted onto slides, and air-dried overnight at 4°C. The sections were treated with concentrated ammonia water for 10 minutes, rinsed thoroughly with purified water, mounted with glycerol gelatin, and scanned using a digital slide scanner (3DHISTECH, Pannoramic MIDI). Dendritic spine density was quantified using ImageJ v1.54p.

#### TUNEL staining

The sections were stained with a TUNEL apoptosis detection kit (G1504, Servicebio) according to the manufacturer’s instructions. Briefly, the sections were incubated with proteinase K working solution at 37°C for 20 minutes, followed by incubation with 0.5% Triton X-100 at room temperature for 20 minutes. A mixture of TdT enzyme, dUTP, and buffer at a ratio of 2:5:50 was applied to cover the tissue, and the sections were incubated at 37°C for 1 hour. DAPI solution was added and incubated at room temperature for 10 minutes in the dark. After washed with PBS, the sections were mounted with anti-fade mounting medium. The sections were scanned using a digital slide scanner (3DHISTECH, Pannoramic MIDI). The density of TUNEL-positive apoptotic cells was quantified using ImageJ v1.54p.

### Enzyme-linked immunosorbent assay (ELISA)

The lungs and hippocampus were homogenized in an appropriate volume of phosphate-free lysis buffer containing protease and phosphatase inhibitors. The homogenates were centrifuged at 12,000 rpm for 15 minutes at 4°C. The total protein concentration in the samples was determined using a BCA kit according to the manufacturer’s instructions. The levels of S1P in serum, lungs, and the hippocampus, as well as the levels of cytokines IL-1β, IL-6, TNF-α, CCL2, IL-4, and IL-10 in the hippocampus, were measured using ELISA kits following the manufacturers’ protocols. The concentrations in tissues were normalized to total protein content.

### Transcriptome sequencing and bioinformatics analysis

#### RNA extraction and quality control

Approximately 50–100 mg of lung or hippocampal tissue was homogenized using TRIzol reagent and a tissue grinder. Total RNA was extracted following the standard phenol-chloroform method. The RNA concentration and purity (OD260/280 and OD260/230 ratios) were measured using a NanoDrop 2000 spectrophotometer (Thermo Fisher Scientific, USA). RNA integrity was assessed using an Agilent 2100 Bioanalyzer (Agilent Technologies, Santa Clara, CA, USA), and samples with an RNA integrity number (RIN) ⩾ 6.0 were considered acceptable.

#### Library construction and sequencing

High-quality total RNA was used to enrich mRNA with poly-A tails using magnetic beads. The enriched mRNA was fragmented, and the fragmented mRNA was used as a template to synthesize double-stranded cDNA using random primers. The double-stranded cDNA underwent end repair, A-tailing, adapter ligation, and PCR amplification to generate the final cDNA library. Library quality was assessed using the Agilent 2100 Bioanalyzer, and the library concentration was determined by Qubit fluorometric quantification. Sequencing was performed on the NovaSeq 6000 platform (Illumina, San Diego, CA, USA) in paired-end 150 bp (PE150) mode, with a target of at least 6 Gb of clean reads per sample. Transcriptome sequencing was conducted by Shanghai Ouyi Biomedical Technology Co., Ltd.

#### Sequencing data processing

The raw sequencing data (raw reads) were quality-controlled and filtered using fastp software (v0.23.2). Reads containing adapters, low-quality bases (Q < 20), excessively high N ratios (>5%), or lengths less than 50 bp were removed to obtain high-quality clean reads. Using the mouse reference genome (GRCm39, Ensembl version 109) as a reference, the clean reads were aligned to the genome using HISAT2 (v2.2.1) with default parameters. The number of reads mapped to each gene was counted using htseq-count to generate a raw count matrix. Genes with zero counts across all samples were removed.

#### Differential expression analysis

Principal component analysis (PCA) was performed to assess sample reproducibility and inter-group separation. Differential expression analysis was conducted using DESeq2 (v1.38.3). The gene count matrix was normalized to the BaseMean value to estimate expression levels. Fold changes were calculated, and statistical significance was assessed using a negative binomial distribution test. Differentially expressed genes (DEGs) were identified with a threshold of p < 0.05. Volcano plots were generated to visualize the overall distribution of DEGs, and heatmaps were used to display expression differences across samples.

#### Enrichment analysis

The identified DEGs were imported into the clusterProfiler package (v4.6.0) in R. Using the mouse genome as the background, Gene Ontology (GO), Kyoto Encyclopedia of Genes and Genomes (KEGG) pathway, Reactome, and WikiPathway enrichment analyses were performed using DEGs lists. Gene set enrichment analysis (GSEA) was performed on all detected genes in the clusterProfiler package (v4.6.0). The gene set databases included GO, KEGG, Reactome, and WikiPathway. The species was set as *Mus musculus*, with a minimum gene set size of 15 and a maximum gene set size of 500. The weighted enrichment statistic was used to calculate the enrichment score (ES), and the ES was normalized to obtain the normalized enrichment score (NES). The selected GSEA enriched terms were plotted to show the distribution of leading-edge genes.

### Metabolomics and bioinformatics analysis

#### Sample preparation

Approximately 30 mg of lung tissue was mixed with 400 μL of methanol/water (4:1, v/v) containing an internal standard (L-2-chlorophenylalanine, 2 μg/mL) and homogenized using a tissue grinder (45 Hz, 2 min). For serum, 50 μL of sample was mixed with 200 μL of protein precipitation solvent (methanol-acetonitrile, 2:1, v/v) containing an internal standard and vortexed for 1 min. Both the lung and serum samples were then subjected to ultrasonic extraction in an ice-water bath for 10 min, followed by standing at −40°C overnight. The samples were centrifuged at 12,000 rpm for 20 min at 4°C, and 150 μL of the supernatant was transferred into an LC-MS vial with an insert for analysis. All reagents were pre-cooled at 4°C before use.

#### Liquid chromatography-mass spectrometry analysis (LC-MS)

A 5 μL aliquot of the extracted sample was injected into a Waters ACQUITY UPLC I-Class Plus/Thermo QE HF mass spectrometry system. Chromatographic separation was performed on an ACQUITY UPLC HSS T3 column (100 mm × 2.1 mm, 1.8 μm) maintained at 45°C. Mobile phase A consisted of 0.1% formic acid in water, and mobile phase B consisted of 0.1% formic acid in acetonitrile, with a flow rate of 0.35 mL/min. The ion source was HESI. Mass spectra were acquired in both positive and negative ion modes using a DDA (data-dependent acquisition) mode with a full MS/dd-MS² (TOP 10) scan pattern.

#### Data processing

The raw data were processed using the metabolomics software XCMS (v4.6.3) for baseline filtering, peak detection, integration, and retention time correction. Compound identification was performed based on retention time (RT), accurate mass, MS/MS fragments, and isotopic distribution using The Human Metabolome Database (HMDB), Lipidmaps (v2.3), METLIN, and the LuMet-Animal local database. The qualitative scoring system had a maximum of 80 points: 20 points for accurate precursor mass matching, 20 points for MS/MS fragment matching, 20 points for isotopic distribution matching, and 20 points for retention time matching. Compounds with a score < 36 were considered inaccurately identified and were removed. After filtering, positive and negative ion data were combined. Ion peaks with a QC sample RSD > 0.3 were removed. Features with >50% zero values within any group were eliminated; the remaining zero values were replaced with half of the minimum ion intensity across all samples. All the data were then log₂-transformed to generate the final data matrix for subsequent analysis. Metabolomics was performed by Shanghai Ouyi Biomedical Technology Co., Ltd.

#### Differentially abundant metabolite analysis

Unsupervised principal component analysis (PCA) was used to assess the overall sample distribution and analytical stability. Supervised orthogonal partial least-squares discriminant analysis (OPLS-DA) was then performed to distinguish global metabolic differences between groups, and variable importance in projection (VIP) values were calculated for each metabolite to evaluate its contribution to inter-group differences. A t-test was used to determine the significance of differences between groups. Metabolites with a VIP ⩾ 1.0 and p < 0.05 were considered differentially abundant. Volcano plots were generated to visualize the overall distribution of differentially abundant metabolites, and heatmaps were used to display expression differences across samples.

#### Enrichment analysis

The list of differentially abundant metabolites was imported into MetaboAnalyst 6.0 (https://www.metaboanalyst.ca) for metabolic pathway enrichment analysis. The species was set as *Mus musculus*, and the KEGG database was used. Hypergeometric testing was applied to calculate enrichment significance, with p < 0.05 as the threshold for significantly enriched pathways. GSEA was performed on all identified metabolites using the in the clusterProfiler package (v4.6.0). The KEGG database was used, with the species set to *Mus musculus*. The minimum metabolite set size was set to 5 and the maximum was set to 200. The weighted enrichment statistic was used to calculate the enrichment score (ES), and the ES was normalized to obtain the normalized enrichment score (NES). The selected GSEA pathways were plotted to show the distribution of leading-edge metabolites.

#### Quantitative real-time PCR (qPCR)

Total RNA was reverse transcribed using the All-In-One 5X RT MasterMix Kit (G592, Applied Biological Materials Inc.). The reverse transcription reaction (20 μL) contained 2 μg of total RNA, 4 μL of 5× RT MasterMix, and RNase-free ddH₂O to bring the volume to 20 μL. The reaction conditions were as follows: 37°C for 15 minutes (reverse transcription), followed by 85°C for 5 seconds (enzyme inactivation). The resulting cDNA was used for qPCR. Amplification was performed on a Jena real-time PCR system using the BlasTaq™ 2X qPCR MasterMix Kit (G891, Applied Biological Materials Inc.). The qPCR mixture (10 μL) consisted of 5 μL of TB Green Premix Ex Taq II (2×), 0.4 μL of forward primer (10 μM), 0.4 μL of reverse primer (10 μM), 0.4 μL of ROX Reference Dye (50×), 1 μL of cDNA template (diluted 5–10-fold), and RNase-free ddH₂O to a final volume of 10 μL. The thermal cycling protocol was as follows: initial denaturation at 95°C for 30 seconds; 40 cycles of denaturation at 95°C for 5 seconds and annealing/extension at 60°C for 30 seconds; followed by a melt curve analysis (95°C for 15 seconds, 60°C for 1 minute, and 95°C for 15 seconds) to verify amplification specificity. The sequences of primers used in the experiments were listed in Supplementary Material 2. GAPDH was employed as the reference gene. The values of the control group was normalized to 1.

### Western blotting

Lung or hippocampal tissues were minced and placed into pre-chilled grinding tubes. Ten volumes of RIPA lysis buffer (Beyotime, Shanghai, China) containing 1% protease inhibitor cocktail and 1% phosphatase inhibitor were added. The tissues were homogenized using a tissue grinder at 60 Hz for 90 seconds, then lysed on ice for 30 minutes with vortexing every 10 minutes. The homogenate was centrifuged at 12,000 rpm for 15 minutes at 4°C, and the supernatant (total protein extract) was transferred to a new EP tube. Protein concentrations were determined using a BCA protein assay kit. All samples were diluted to the same concentration with RIPA lysis buffer, mixed with 5× SDS loading buffer containing β-mercaptoethanol, and denatured by heating at 100°C for 10 minutes in a metal bath. Proteins were separated by SDS-PAGE with 20 μg of total protein loaded per well. After electrophoresis, the target proteins were transferred onto polyvinylidene difluoride (PVDF) membranes (Merck Millipore, Billerica, MA, USA). The membranes were blocked with TBST containing 5% non-fat dry milk for 1 hour at room temperature, followed by overnight incubation at 4°C with diluted primary antibodies. After washing, the membranes were incubated with HRP-conjugated secondary antibodies for 1 hour at room temperature. The protein bands were visualized using an enhanced chemiluminescence (ECL) kit and imaged with a chemiluminescence imaging system (GE Amersham Imager 800, GE Healthcare Life Sciences). Band intensities were quantified using ImageJ v1.53. The relative expression level of each target protein was normalized to that of GAPDH, and that of the control group was normalized to 1.

### Statistical analysis

The statistician was blinded to the group assignment of the samples. Statistical analyses were conducted using GraphPad Prism software v9.5. The data were presented as the mean ± standard error of the mean (SEM). For comparisons between two groups, an unpaired t-test was used. For multiple group comparisons, one-way ANOVA followed by Tukey’s multiple comparisons test was employed. A p-value < 0.05 was considered statistically significant, and asterisks were used to indicate significance in the Figure.

## Results

### Intranasal instillation of bleomycin induces pulmonary fibrosis and concomitant emotion-like disorders in mice

Numerous clinical studies have reported a high incidence of mood disorders in patients with IPF [4–6]. To determine whether a similar phenomenon occurs in an animal model and to explore the underlying mechanisms, we established an IPF mouse model by a single intratracheal instillation of bleomycin. The experimental procedure is shown in Figure 1A. At 28 days after bleomycin instillation, H&E and Masson staining revealed thickening of the bronchial and alveolar walls, disruption of the alveolar architecture (Figure 1B), and a marked increase in the collagen content (Figure 1C) compared with those of the control group. These data confirmed the successful establishment of the IPF animal model. We evaluated animal mood behavior using the OFT, SPT, TST, and FST. The results showed that IPF mice spent significantly less time in the central area of the open field (Figure 1D, 1E), exhibited reduced sucrose preference (Figure 1F), and showed increased immobility time in both the TST and FST (Figure 1G, 1H). These findings demonstrate that IPF mice display emotional disorders characterized by anxiety, anhedonia, and despair-like behavior.

**Fig. 1.**
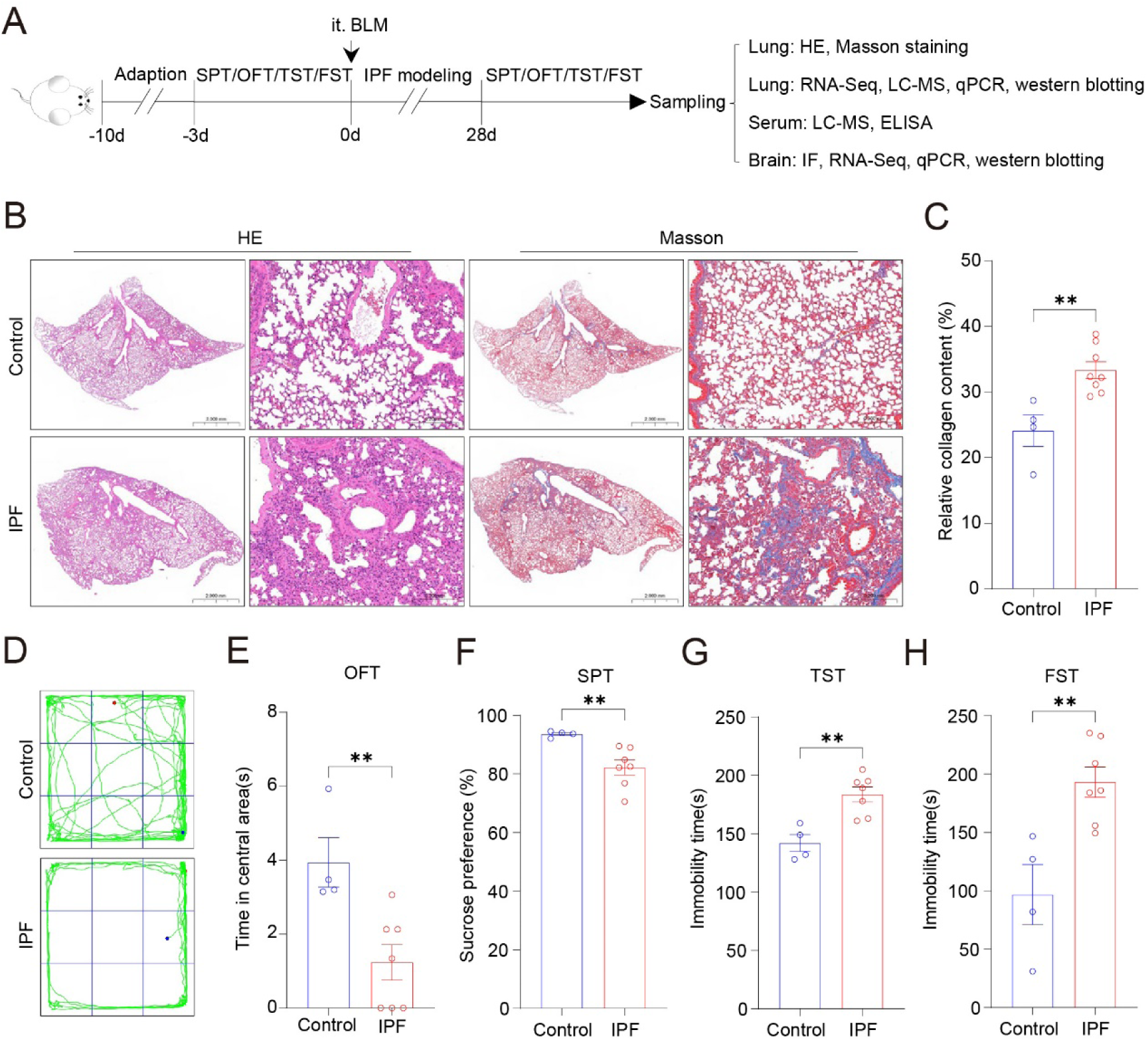
IPF mice exhibit emotional disorders. (A) Schematic diagram of the study design. (B) Representative images of H&E and Masson staining of lung tissue. Bar = 2 mm in the panorama images and 0.2 mm in zoom images. (C) Collagen content in lung tissue. (D) Representative trace plots of the open field test (OFT). (E) Time spent in the central area of the open field. (F) Percentage of sucrose consumption in the sucrose preference test (SPT). (G) Immobility time in the tail suspension test (TST). (H) Immobility time in the forced swimming test (FST). Control: n=4, IPF: n=7. ** *p* < 0.01.

### Pulmonary fibrosis leads to synaptic damage and cell death in the hippocampus

The hippocampus is a critical brain region involved in emotional regulation. To investigate hippocampal changes in IPF mice, we examined the expression of the synapse-associated proteins PSD95 and SYP using immunofluorescence (Figure 2A, 2B). Compared with the control group, IPF mice presented significantly lower fluorescence intensities of PSD95 and SYP in the CA1, CA3, and DG subregions of the hippocampus (Figure 2C–2H), indicating synaptic dysfunction. Next, Golgi staining was performed to visualize hippocampal dendritic spines, and their density was quantified. IPF mice showed a marked reduction in dendritic spine density in the hippocampus relative to controls (Figure 2I, 2K). Furthermore, the result from TUNEL staining revealed that the density of TUNEL-positive cells was significantly higher in the CA1, CA3, and DG regions of IPF mice than those of the control mice (Figure 2J, 2L), suggesting increased cell death in the hippocampus. These findings indicate that pulmonary fibrosis disrupts hippocampal synaptic function and induces cell death, thereby driving mood disorders.

**Fig. 2.**
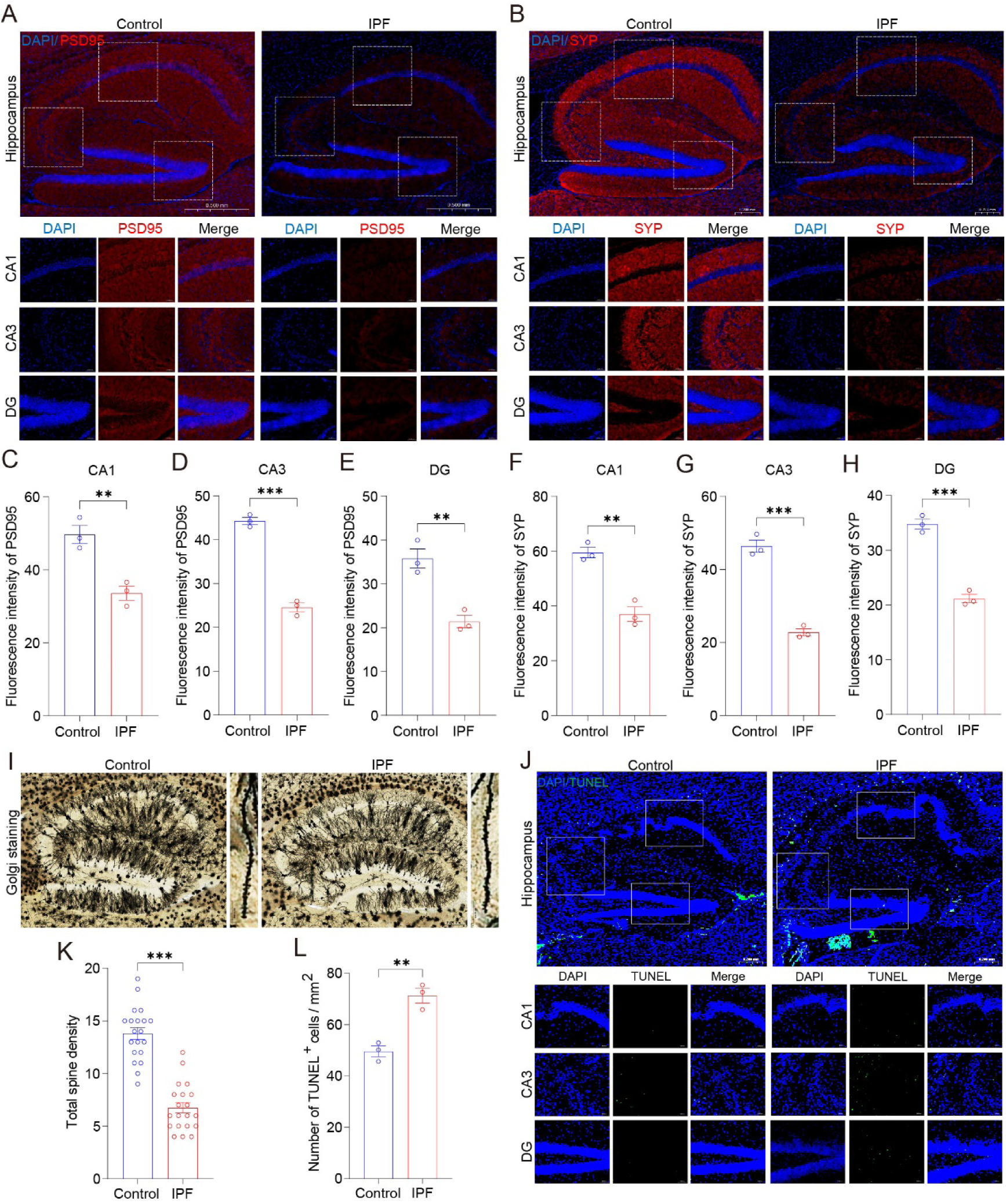
Hippocampal synaptic function is impaired and cell death is increased in IPF mice. (A) Immunofluorescence staining of PSD95 in the hippocampus. Bar = 0.5 mm in panorama images and 0.05 mm in zoom images. (B) Immunofluorescence staining of SYP in the hippocampus. Bar = 0.2 mm in panorama images and 0.05 mm in zoom images. (C–E) Fluorescence intensity of PSD95 in the CA1, CA3, and DG subregions of the hippocampus. (F–H) Fluorescence intensity of SYP in the CA1, CA3, and DG subregions of the hippocampus. (I) Golgi staining and representative dendritic spines in the hippocampus. Bar = 0.2 mm in panorama images and 0.01 mm in zoom images. (J) TUNEL staining in the hippocampus. Bar = 0.2 mm in panorama images and 0.05 mm in zoom images. (K) Dendritic spine density in the hippocampus as assessed by Golgi staining. (L) Density of TUNEL-positive cells in the hippocampus. Control: n = 3, IPF: n = 3. ** *p* < 0.01, \*\*\**p* < 0.001.

### Hippocampal microglial and astrocytic activation and neuroinflammation in mice with pulmonary fibrosis

Glial cells and glia-mediated neuroinflammation play important roles in the central nervous system. We therefore assessed glial activation by immunofluorescence staining for the microglial marker IBA1 and the astrocytic marker GFAP (Figure 3A, 3G). Compared with the control mice, IPF mice exhibited significantly increased fluorescence intensity and density of IBA1-positive cells in the hippocampus (Figure 3B, 3C). The total branch length and number of branch points of IBA1-positive cells were markedly reduced (Figure 3D, 3E). The branching complexity of IBA1-positive cells decreased, revealed by Sholl analysis (Figure 3F). Similarly, the fluorescence intensity and density of GFAP-positive cells in the hippocampus were significantly increased (Figure 3H, 3I), accompanied by marked reductions in total branch length and branch point number (Figure 3J, 3K) and decreased branching complexity (Figure 3F). These data indicate that microglia and astrocytes are activated in the hippocampus of IPF mice. Furthermore, we measured hippocampal inflammatory cytokine levels via ELISA. Compared with controls, IPF mice presented elevated levels of the proinflammatory cytokines IL-1β, IL-6, TNF-α, and CCL2, and decreased levels of the anti-inflammatory cytokines IL-4 and IL-10 in the hippocampus, indicating the presence of neuroinflammation. These findings suggest that pulmonary fibrosis induces glial activation and neuroinflammation in the hippocampus.

**Fig. 3.**
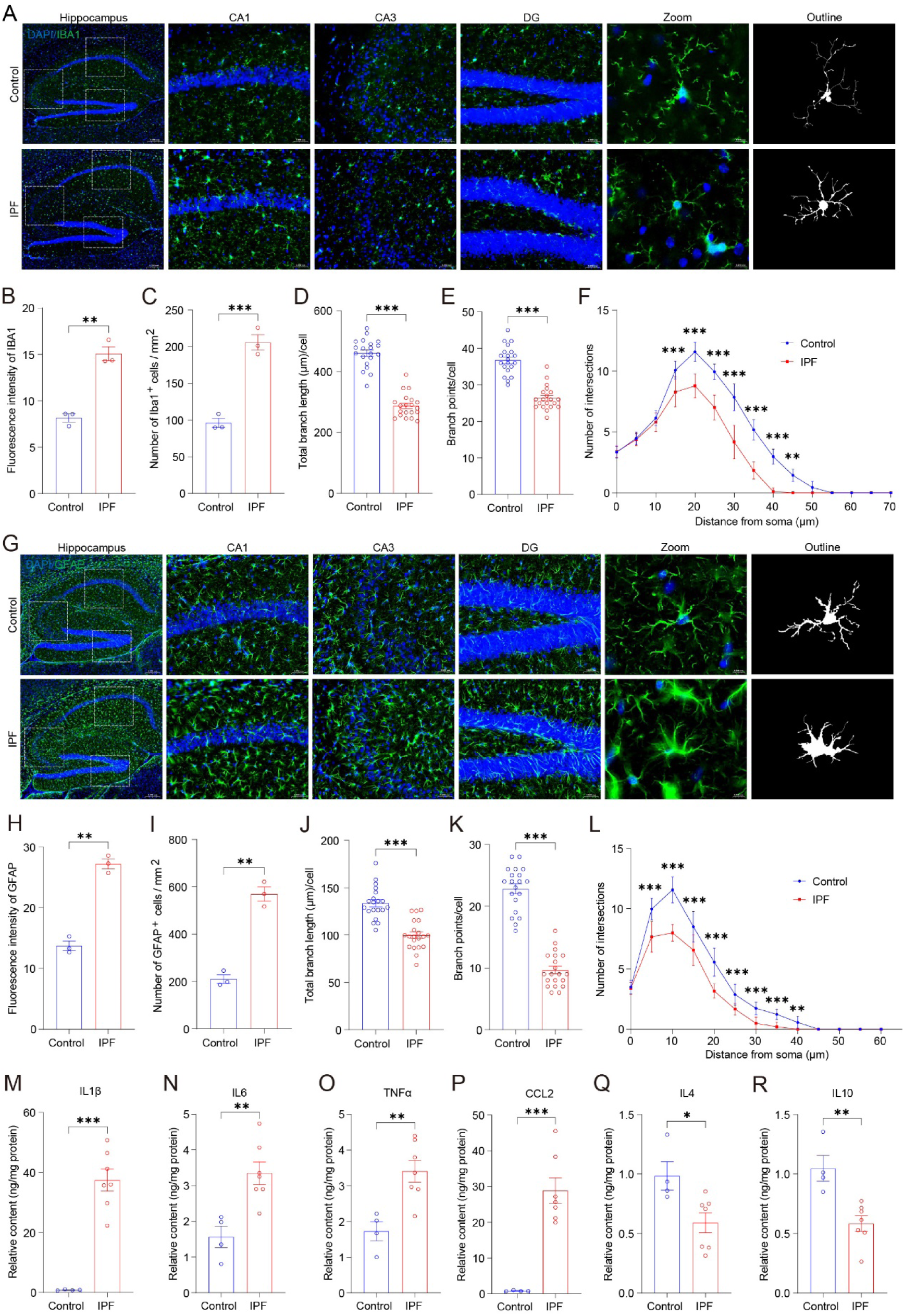
Microglial and astrocytic activation and neuroinflammation in the hippocampus of IPF mice. (A) Immunofluorescence staining of IBA1 and morphological analysis of microglia in the hippocampus. Bar = 0.2 mm in panorama images, 0.05 mm in CA1, CA3 and DG zoom inages, and 0.01 mm in single cell (outline) zoom images. (B) Fluorescence intensity of IBA1 in the hippocampus. (C) Density of IBA1-positive cells. (D) Total branch length of microglia. (E) Number of branch points of microglia. (F) Quantification of microglial branch complexity by Sholl analysis. (G) Immunofluorescence staining of GFAP and morphological analysis of astrocytes in the hippocampus. Bar = 0.2 mm in panorama images, 0.05 mm in CA1, CA3 and DG zoom images, and 0.01 mm in single cell (outline) zoom images. (H) Fluorescence intensity of GFAP in the hippocampus. (I) Density of GFAP-positive cells. (J) Total branch length of astrocytes. (K) Number of branch points of astrocytes. (L) Quantification of astrocytic branch complexity by Sholl analysis. (M–R) Relative levels of the cytokines IL-1β, IL-6, TNF-α, CCL2, IL-4, and IL-10 in the hippocampus. Immunofluorescence: Control n = 3, IPF n = 3; ELISA: Control n = 4, IPF: n = 7. ** *p* < 0.01, \*\*\**p* < 0.001.

### Upregulation of sphingolipid metabolism in fibrotic lungs

To identify pulmonary changes and factors contributing to mood disorders, we collected lung tissues from IPF and control mice and performed transcriptomic sequencing followed by bioinformatics analysis. Principal component analysis (PCA) revealed a clear separation between the transcriptomes of IPF and control lungs (Figure 4A), indicating substantial differences. Compared with the control group, we identified 593 downregulated genes and 390 upregulated genes in IPF lung tissues (fold change > 1.5, p < 0.05; Figure 4B, 4C). Enrichment analysis of the upregulated genes showed significant enrichment of biological processes and pathways related to sphingolipid metabolism (Figure 4D). GSEA of all genes further confirmed the enrichment of lung fibrosis and sphingolipid metabolism pathways (Figure 4E–4G). These results suggest that sphingolipid metabolism is altered in fibrotic lung tissue.

**Fig. 4.**
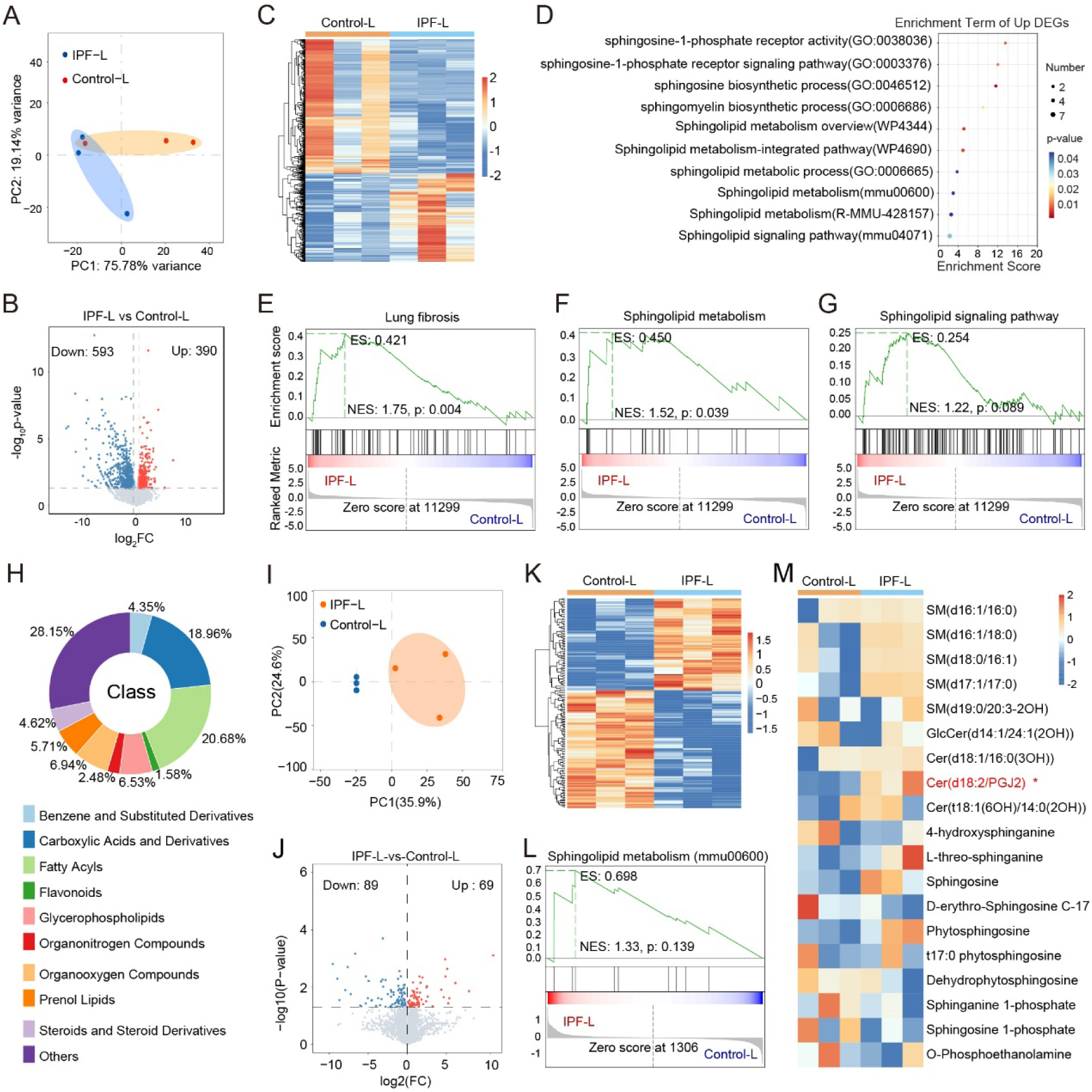
Transcriptomic and metabolomic analyses reveal upregulated sphingolipid metabolism in the lungs of IPF mice. (A) PCA of transcriptomic profiles in lung tissues of IPF mice. (B) Volcano plot of differentially expressed genes in lung tissues. (C) Heatmap of differentially expressed genes in the lungs. (D) Enrichment analysis of differentially expressed genes in lung tissues. (E–G) GSEA of lung transcriptomic data related to lung fibrosis, sphingolipid metabolism, and GPCR signaling pathways, respectively. (H) Classification of lung tissue metabolites identified by LC-MS. (I) PCA of metabolomic profiles. (J) Volcano plot of differentially abundant metabolites. (K) Heatmap of differentially abundant metabolites. (L) GSEA of metabolomic data related to sphingolipid metabolism. (M) Heatmap of metabolites involved in sphingolipid metabolism in lung tissues.

Since sphingolipids are a class of chemical molecules, we next performed LC-MS-based metabolomic analysis of lung tissues to identify specific molecular changes. The most abundant metabolite classes in mouse lung tissues were carboxylic acids and derivatives, and fatty acyls (Figure 4H). PCA clearly separated the metabolomes of IPF and control lungs (Figure 4I), indicating significant metabolic differences. Compared with the control, 89 metabolites were downregulated and 69 were upregulated in IPF lung tissues (p < 0.05; Figure 4J, 4K). GSEA of the differential metabolites indicated an upregulation of sphingolipid metabolism, albeit not statistically significant (Figure 4L). Based on literature reports of molecules involved in sphingolipid metabolism [30, 31], we selected these molecules and displayed their relative abundances and differences via a heatmap. The results showed that only Cer(d18:2/PGJ2) was significantly elevated in IPF mouse lung tissues (Figure 4M). These findings indicate that sphingolipid metabolism is altered in IPF lung tissues and may contribute to the development of anxiety- and depression-like behaviors.

### Elevated serum S1P in IPF mice drives emotional disorders

Metabolites derived from lung tissue can act on the central nervous system via the bloodstream. Therefore, we performed LC-MS-based metabolomic analysis of serum samples followed by bioinformatics analysis. The predominant metabolite classes in mouse serum were carboxylic acids and derivatives, fatty acyls, and glycerophospholipids (Figure 5A). PCA revealed a clear separation between the serum metabolomes of IPF and control mice (Figure 5B), indicating significant metabolic differences. Compared with the control group, the serum of IPF mice contained 74 downregulated and 154 upregulated metabolites (p < 0.05; Figure 5C, 5D). We further analyzed the abundance of metabolites involved in sphingolipid metabolism and found that S1P and its derivatives, sphingosine 1-phosphate (d16:1-P) and C17 sphingosine-1-phosphate, were markedly elevated (Figure 5E, 5F). The ELISA results confirmed that the serum S1P levels were significantly higher in IPF mice than in controls (Figure 5G).

**Fig. 5.**
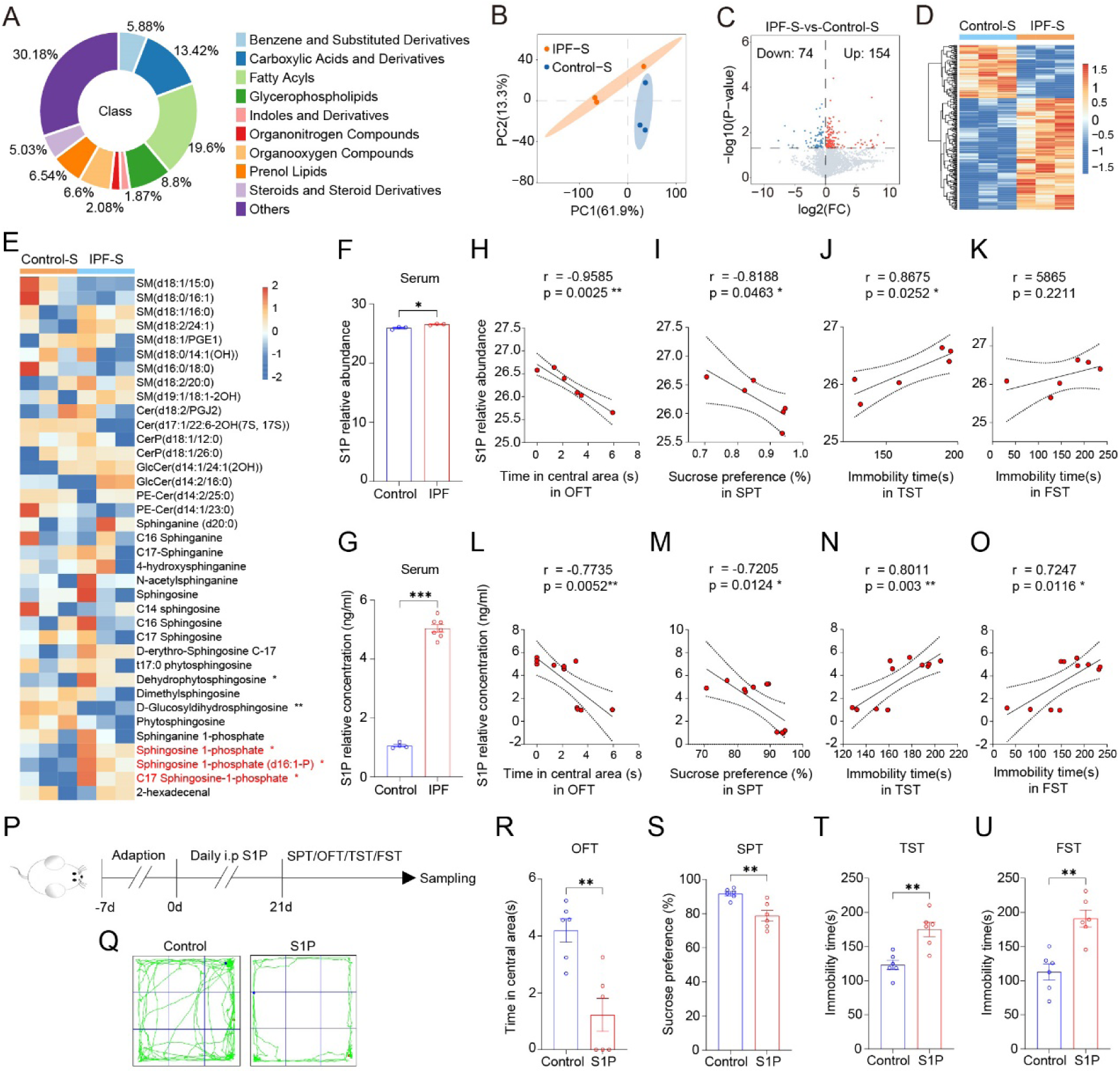
Elevated serum S1P in IPF mice drives emotional disorders. (A) Classification of serum metabolites in IPF mice identified by LC-MS. (B) PCA of metabolomic profiles. (C) Volcano plot of differentially abundant metabolites. (D) Heatmap of differentially abundant metabolites. (E) Heatmap of metabolites involved in sphingolipid metabolism in the serum. (F) Relative abundance of serum S1P measured by LC-MS. (G) Relative concentration of serum S1P measured by ELISA. (H–K) Correlations between relative abundance of serum S1P and time spent in the central area of the OFT, sucrose preference, immobility time in the TST, and immobility time in the FST, respectively. (L–O) Correlations between relative concentration of serum S1P and time spent in the central area of the OFT, sucrose preference, immobility time in the TST, and immobility time in the FST, respectively. (P) Experimental timeline of S1P injection. (Q) Representative trace plots of the OFT. (R) Time spent in the central area of the open field. (S) Percentage of sucrose consumption in the SPT. (T) Immobility time in the TST. (U) Immobility time in the FST. ELISA: Control n = 4, IPF n = 7; Behavior: Control n = 6, S1P: n = 6. * *p* < 0.05, ** *p* < 0.01, \*\*\**p* < 0.001.

On the basis of these findings, we hypothesized that elevated serum S1P in IPF mice drives the onset of mood disorders. We first analyzed the relationships between relative abundance/concentration of S1P in the serum and behavioral outcomes. Pearson correlation analysis revealed that S1P levels were significantly negatively correlated with time spent in the center of the open field and sucrose preference (p < 0.05) and significantly positively correlated with immobility time in the TST and FST (p < 0.05), with the exception of the correlation between S1P abundance and immobility time in the FST (Figure 5H–5O). These results suggest that serum S1P levels are positively associated with the affective symptoms in IPF mice.

To validate this hypothesis, we intraperitoneally injected mice with a supraphysiological dose of 5 mg S1P /kg body weight daily for 21 days and then assessed behavioral changes (Figure 5P). Compared with the control, S1P injection resulted in decreased time spent in the center of the open field, reduced sucrose preference, and increased immobility time in the TST and FST (Figure 5Q–5U), indicating that peripheral administration of a high dose of S1P induces anxiety- and depression-like symptoms. These findings demonstrate that elevated serum S1P levels in IPF mice drive emotion-like disorders.

### Inhibition of Sphk1 reduces peripheral S1P levels and alleviates emotional disorders in IPF mice

Although our findings confirmed that serum S1P induces mood disorders in IPF mice, we sought to determine whether the elevated serum S1P originates from lung tissue. We therefore measured S1P concentrations in lung tissue by ELISA and found no change (Figure 6A). This result, which seemed inconsistent with our transcriptomic and serum ELISA findings (Figure 4M and 5E), prompted us to further investigate this issue. We reviewed the literature and summarized the metabolic pathways and related genes involved in S1P metabolism [30–34], categorizing them according to their functions (Figure 6B, 6C). We then re-examined the expression of S1P metabolism-related genes in the lung transcriptomic data. The data showed upregulation of multiple genes involved in the synthesis of S1P or its precursors, including Sptlc2 (encoding serine palmitoyltransferase, which catalyzes the conversion of serine and palmitoyl-CoA to 3-ketosphinganine), Asah1 (encoding N-acylsphingosine amidohydrolase, which hydrolyzes ceramide to sphingosine), Sgms1 (encoding sphingomyelin synthase, which also hydrolyzes ceramide to sphingosine), and Sphk1 (encoding sphingosine kinase, which phosphorylates sphingosine to S1P) [30–34]. Interestingly, the expression of Mfsd2a and Mfsd2b, which encode S1P exporters, was also significantly upregulated (Figure 6D).

**Fig. 6.**
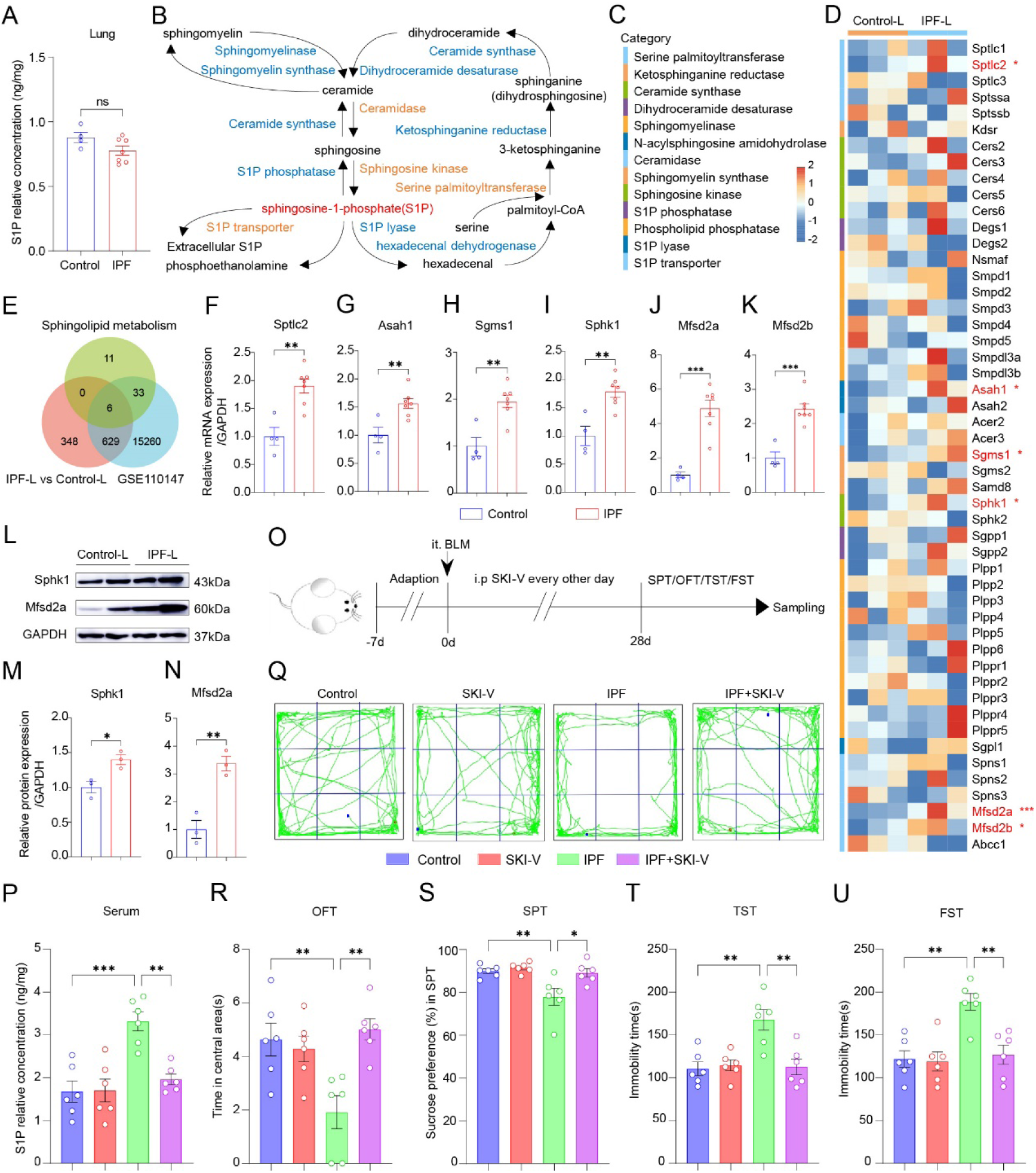
Inhibition of Sphk1 reduces peripheral blood S1P and alleviates emotional disorders in IPF mice. (A) Relative concentration of S1P in lung tissue measured by ELISA. (B) Schematic diagram of S1P metabolic pathways. (C) Classification of S1P metabolic enzymes. (D) Heatmap of S1P metabolism-related gene expression in lung tissue based on transcriptomic sequencing. Red indicates differentially expressed genes. (E) Venn diagram showing the number of common differentially expressed genes related to sphingolipid metabolism identified in the lung transcriptome of IPF mice and in lung tissue microarrays of IPF patients (GSE110147). (F–K) Validation of S1P metabolism-related gene expression in lung tissue by qPCR. (L–N) Validation of Sphk1 and Mfsd2a protein expression in lung tissue by Western blotting. (O) Experimental timeline for SKI-V treatment in IPF mice with comorbid mood disorders. (P) Relative concentration of S1P in the serum measured by ELISA. (Q) Representative trace plots of the OFT. (R) Time spent in the central area of the open field. (S) Percentage of sucrose consumption in the SPT. (T) Immobility time in the TST. (U) Immobility time in the FST. qPCR: Control n = 4, IPF n = 7; WB: Control n = 3, IPF: n = 3; Behavior: Control n = 6, IPF n = 6. ns: not significant. * *p* < 0.05, ** *p* < 0.01, \*\*\**p* < 0.001.

To assess the relevance of our findings to clinical samples, we analyzed transcriptomic data from IPF patients in the GEO database. The analysis revealed that both clinical IPF patients and IPF mice shared common differentially expressed genes related to sphingolipid metabolism, specifically the six genes mentioned above (Figure 6E), indicating that our model recapitulates key gene expression features of human IPF, particularly in sphingolipid metabolism. We next validated the expression of these genes by qPCR, and the results were consistent with the sequencing data (Figure 6F–6K). Furthermore, we examined the protein expression of Sphk1, the rate-limiting enzyme for S1P synthesis, and Mfsd2a, the S1P exporter, by Western blotting. The results showed that both Sphk1 and Mfsd2a protein levels were significantly elevated in IPF mice compared with controls (Figure 6L-6N). These findings suggest that although S1P synthesis is increased in fibrotic lung tissue, S1P export from cells is also enhanced, which likely explains why S1P levels remain unchanged in lung tissue but are elevated in the serum of IPF mice. These data support our hypothesis that the elevated serum S1P in IPF mice originates from lung tissue.

To investigate the impact of pulmonary S1P metabolism on emotional behavior, we administered the Sphk1 inhibitor SKI-V by daily intraperitoneal injection to mice following intranasal instillation of bleomycin (Figure 6O). After 28 days, the serum S1P level was significantly lower in SKI-V-treated mice than that in IPF mice (Figure 6P), further confirming that the elevated serum S1P in IPF mice originates from the fibrotic lungs. Compared with the IPF group, the SKI-V-treated group presented increased time spent in the center of the open field, increased sucrose preference, and decreased immobility time in the TST and FST, indicating that SKI-V alleviated anxiety- and depression-like symptoms in IPF mice (Figure 6Q–6U). These findings demonstrate that inhibiting Sphk1 reduces S1P synthesis in fibrotic lungs and lowers serum S1P levels, thereby improving mood disorders in IPF model.

### Peripherally derived S1P in IPF model activates the hippocampal S1PR1/PI3K/PKA/CREB signaling pathway

To investigate how IPF affects hippocampal function and thereby drives mood disorders, we performed transcriptomic sequencing and bioinformatics analysis on hippocampal tissues from IPF and control mice. Principal component analysis (PCA) revealed a clear separation between the hippocampal transcriptomes of IPF and control mice (Figure 7A), indicating substantial differences. Compared with the control, 286 genes were downregulated and 461 genes were upregulated in the hippocampus of IPF mice (p < 0.05; Figure 7B, 7C). Interestingly, enrichment analysis of the upregulated genes revealed that multiple enriched terms were associated with sphingolipid binding, S1P receptor activation, and GPCR (S1P receptor [35]) signaling (Figure 7D). GSEA using all the hippocampal genes further demonstrated upregulation of the GPCR pathway in IPF mice compared with controls (Figure 7E). Taken together, these results suggest that the S1P receptor (S1PR) may be activated in the hippocampus of IPF mice. Thus, We next analyzed the expression of the five S1PR subtypes in the hippocampus [35, 36]. Transcriptomic and qPCR data showed that among the five subtypes, only S1PR1 was significantly upregulated in the hippocampus of IPF mice relative to the control (Figure 7F, 7G), which was confirmed by Western blotting (Figure 7H, 7I). These findings indicate that the activation of S1PR1 in the hippocampus of IPF mice may contribute to the development of emotional disorders.

**Fig. 7.**
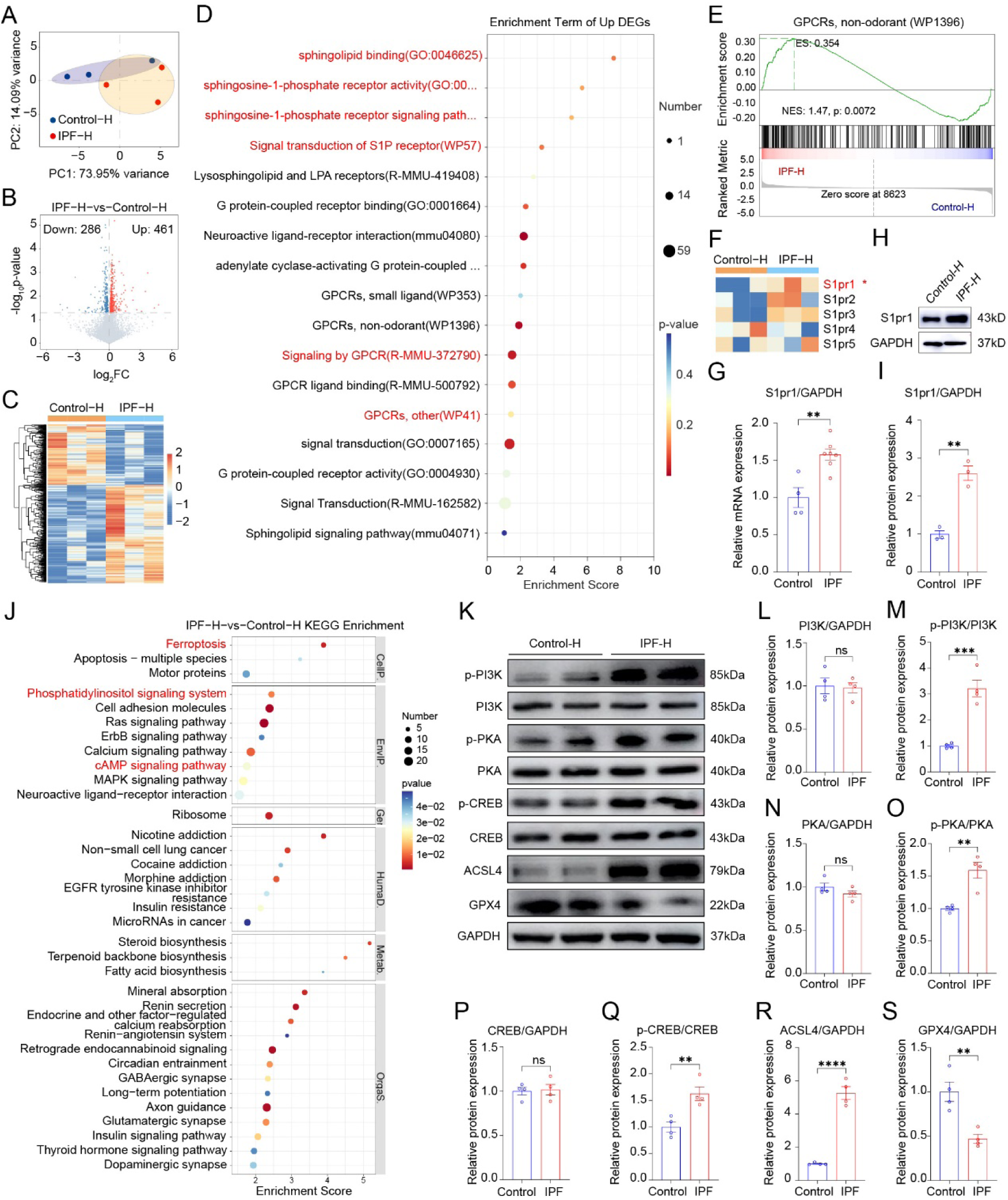
Hippocampal S1PR1 is activated and regulates ferroptosis via the PI3K/PKA/CREB signaling pathway in IPF mice. (A) PCA of transcriptomic profiles in the hippocampus of IPF mice. (B) Volcano plot of the differentially expressed genes. (C) Heatmap of differentially expressed genes. (D) Enrichment analysis of upregulated differentially expressed genes. (E) GSEA of hippocampal transcriptomic data related to the GPCR signaling pathway. (F) Heatmap of S1P receptor expression levels. (G) Validation of S1pr1 expression in the hippocampus by qPCR. Control: n = 4, IPF: n = 7. (H, I) Validation of S1PR1 protein expression in the hippocampus by Western blotting. Control: n = 3, IPF: n = 3. (J) KEGG enrichment analysis of differentially expressed genes. (K–S) Validation of p-PI3K, PI3K, p-PKA, PKA, p-CREB, CREB, ACSL4, and GPX4 protein expression in the hippocampus by Western blotting. Control: n = 6, IPF: n = 6. ns: not significant. ** *p* < 0.01, \*\*\**p* < 0.001, **** *p* < 0.0001.

Because S1P is widely present in the nervous system, including the hippocampus, we sought to determine whether S1PR1 in the hippocampus of IPF mice is activated by local S1P. Therefore, we measured hippocampal S1P levels via ELISA. The results showed that, compared with controls, S1P levels in the hippocampus of IPF mice were not elevated but rather slightly decreased, although the difference was not statistically significant (Figure S1A). Bioinformatics analysis revealed that the downregulated genes in the hippocampus of IPF model were enriched in sphingolipid metabolism (Figure S1B). GSEA also showed a trend toward downregulation of sphingolipid metabolism-related pathways, albeit not significant (Figure S1C). Transcriptomic and qPCR data indicated that the expression of S1P metabolism- and transport-related genes, including Sphk1 and Mfsd2a, presented no change in the hippocampus of IPF mice compared with controls, with the exception of Cers2, which was significantly downregulated (Figure S1D–S1I). Together with previous findings, these data suggest that S1PR1 in the hippocampus of IPF mice is activated by serum-derived S1P rather than by local hippocampal S1P.

We next investigated which downstream signaling pathways and biological processes are affected by S1PR1 activation. KEGG enrichment analysis of the DEGs in the hippocampus of IPF animals showed significant enrichment of ferroptosis, the phosphatidylinositol signaling system, and the cAMP signaling pathway (Figure 7J). We examined key molecules in these pathways and biological processes by Western blotting, including p-PI3K, PI3K, p-PKA, PKA, p-CREB, CREB, ACSL4, and GPX4 (Figure 7K). The results showed that, compared with controls, total PI3K, PKA, and CREB levels in the hippocampus of IPF mice were unchanged, whereas p-PI3K, p-PKA, and p-CREB levels were significantly increased (Figure 7L–7Q). The ferroptosis-promoting protein ACSL4 was markedly upregulated (Figure 7R), while the ferroptosis-inhibitory protein GPX4 was significantly downregulated (Figure 7S). Taken together, these findings suggest that serum-derived S1P in IPF mice acts on hippocampal S1PR1 to regulate the PI3K/PKA/CREB signaling pathway and induce ferroptosis.

### Fingolimod ameliorates emotional disorders by alleviating hippocampal neuroinflammation and cell death via the PI3K/PKA/CREB pathway

Given that S1P activates hippocampal S1PR1 and drives mood disorders, we hypothesized that inhibiting S1PR1 function could alleviate IPF-induced mood disorders. Therefore, while establishing the IPF model by bleomycin (BLM) instillation, we treated mice with Fingolimod (FTY720, abbreviated as FTY), a selective functional inhibitor of S1PR1 that is clinically used for the treatment to multiple sclerosis (Figure 8A). Compared with the IPF group, daily intraperitoneal injection of FTY increased the time spent in the center of the open field and sucrose preference, and decreased immobility time in the TST and FST (Figure 8B–8F). Immunofluorescence results revealed that FTY treatment reduced IBA1-positive, GFAP-positive, and TUNEL-positive signals in the hippocampus of IPF mice. The data from ELISA showed that FTY decreased the hippocampal levels of the proinflammatory cytokines IL-1β, IL-6, TNF-α, and CCL2, and increased the levels of the anti-inflammatory cytokines IL-4 and IL-10 in IPF mice. Furthermore, FTY restored the elevated levels of p-PI3K, p-PKA, p-CREB, and ACSL4, as well as the reduced level of GPX4, in the hippocampus of IPF mice. These data indicate that Fingolimod ameliorates anxiety-and depression-like behaviors by alleviating hippocampal neuroinflammation and ferroptosis via the PI3K/PKA/CREB pathway in IPF mice.

**Fig. 8.**
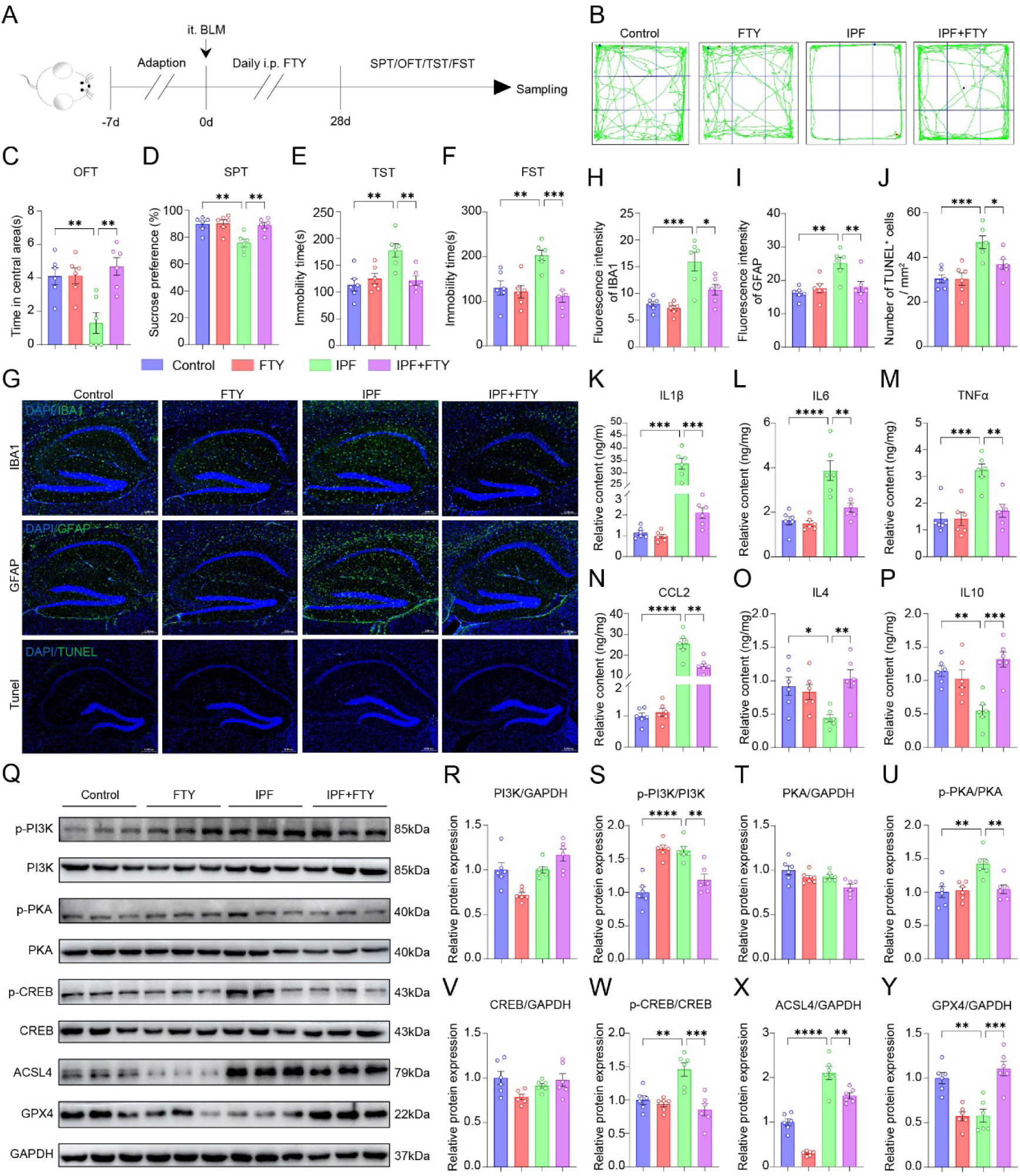
Fingolimod ameliorates emotional disorders in IPF mice by reducing hippocampal neuroinflammation and ferroptosis via the PI3K/PKA/CREB pathway. (A) Experimental timeline of Fingolimod treatment in IPF mice with mood disorders. (B) Representative trace plots of the OFT. (C) Time spent in the central area of the open field. (D) Percentage of sucrose consumption in the SPT. (E) Immobility time in the TST. (F) Immobility time in the FST. (G) Immunofluorescence staining of IBA1, GFAP, and TUNEL in the hippocampus. (H) Fluorescence intensity of IBA1. (I) Fluorescence intensity of GFAP. (J) Density of TUNEL-positive cells. (K–P) Relative levels of the cytokines IL-1β, IL-6, TNF-α, CCL2, IL-4, and IL-10 in the hippocampus. (Q–Y) Protein expression of p-PI3K, PI3K, p-PKA, PKA, p-CREB, CREB, ACSL4, and GPX4 in the hippocampus was assessed by Western blotting. Bar = 0.2 mm, n = 6 for all groups. ns: not significant. ** *p* < 0.01, \*\*\**p* < 0.001, **** *p* < 0.0001.

## Discussion

Although the higher prevalence of depression and anxiety in patients with IPF than in the general population is well documented, the underlying biological mechanisms have remained largely unexplored [3–6]. In this study, we addressed this important yet unresolved question by proposing and validating an original scientific hypothesis: sphingosine-1-phosphate (S1P) derived from fibrotic lungs enters the bloodstream, acts on the S1P receptor S1PR1 in the hippocampus, and subsequently activates the PI3K/PKA/CREB signaling pathway, driving neuroinflammation and ferroptosis in the hippocampus, ultimately leading to the development of emotional disorders. Our findings not only confirm the presence of affective disorders in IPF model but also reveal a molecular pathway from pulmonary fibrosis to brain dysfunction, providing a novel mechanistic explanation and suggesting potential therapeutic strategies for IPF-associated emotional disorders.

S1P is a bioactive sphingolipid that plays a critical regulatory role in various physiological and pathological processes, including cell proliferation, survival, migration, and immune cell trafficking [37]. Under physiological conditions, S1P in the blood is produced primarily by red blood cells and vascular endothelial cells and maintained at relatively stable levels; however, under pathological conditions, S1P production and release can be significantly altered [38, 39]. Previous studies have shown that pulmonary lesions lead to elevated S1P levels, which may serve as a therapeutic target for respiratory diseases [40, 41]. An important discovery of the present study is that fibrotic lungs may be an aberrant source of S1P, which remotely regulates hippocampal function via the bloodstream. This finding links two seemingly independent processes—metabolic changes in pulmonary fibrosis and central mood disorders. Notably, our results echo previous reports on the role of S1P in mood disorders. Studies have suggested that anxiety-like behaviors in animal models are associated with serum S1P levels [42–44]. The unique contribution of this study lies in revealing, for the first time, the role of this signaling molecule in the context of the lung–brain axis: S1P not only acts as a signaling molecule generated within the nervous system but also serves as a long-range signal originating from peripheral organs (particularly the fibrotic lung), crossing the blood–brain barrier or influencing the functional state of the central nervous system through indirect mechanisms. This finding provides a new perspective—from a metabolic standpoint—for understanding the neuropsychiatric complications associated with chronic lung disease.

In our study of the IPF mouse model, we observed a seemingly paradoxical yet biologically significant phenomenon: in fibrotic lung tissue, the expression of the S1P-synthesizing enzyme Sphk1 was markedly increased, whereas the expression of the S1P efflux-associated protein Mfsd2a was also upregulated; however, the local S1P concentration in the lung remained unchanged, whereas blood S1P levels were significantly elevated. This phenomenon suggests that, in the IPF state, the regulation of S1P in lung tissue does not simply lead to accumulation but instead establishes a dynamic pattern of “increased synthesis and increased efflux”. Previous studies have confirmed that Sphk1 is significantly upregulated in both IPF patients and bleomycin-induced pulmonary fibrosis mouse models, and that Sphk1 expression levels are negatively correlated with lung function and positively correlated with fibrosis markers in patients [45, 46]. S1P produced by Sphk1 promotes fibroblast activation and directly drives the development and progression of pulmonary fibrosis by activating the Hippo/YAP signaling pathway and generating mitochondrial reactive oxygen species [47]. Sphk1 knockout or pharmacological inhibition significantly alleviates BLM-induced pulmonary fibrosis and reduces mortality in mice [48]. However, elevated Sphk1 does not necessarily lead to local S1P accumulation. We noted that the upregulation of Sphk1 was accompanied by increased expression of the S1P efflux protein Mfsd2a. Mfsd2a is a sodium-dependent lysophosphatidylcholine (LPC) transporter abundantly expressed on the apical membrane of alveolar type II (AT2) cells. Its functions have recently been extended to include the maintenance of lipid metabolic homeostasis and the regulation of fibrosis. It has been reported that LPC promotes AT2 cell senescence and Drp1-mediated mitochondrial fission via an Mfsd2a-dependent pathway, thereby driving fibrosis following lung injury [49]. Mfsd2a also plays an important role in IPF-associated metabolic reprogramming, and its upregulation may directly participate in the regulation of lipid metabolism disorders during fibrosis [50]. In cystic fibrosis, transcriptional upregulation of Sphk1 is closely associated with abnormal S1P accumulation, and a functional deficiency of the efflux protein Spns2 further exacerbates intracellular S1P retention [51]. In contrast, our study revealed that in IPF, the upregulation of Mfsd2a may promote S1P efflux from the lung, thereby increasing S1P output into the circulation and leading to elevated blood S1P levels. Once S1P enters the bloodstream, it can cross the blood-brain barrier or bind to its receptors on brain microvascular endothelial cells, thereby regulating brain activity. This coordinated upregulation of synthesis and efflux may constitute an important molecular basis for understanding lung–brain axis signaling. More importantly, we found that inhibiting Sphk1 with SKI-V reduced serum S1P levels in IPF mice and alleviated mood symptoms. These findings demonstrate the role of lung-derived S1P in emotional disorders.

Among the five S1P receptors, S1PR1, S1PR2, S1PR3, and S1PR5 are expressed to varying degrees on neurons, astrocytes, oligodendrocytes, microglia and endothelial cells in the central nervous system, whereas S1PR4 is expressed in neurons [52, 53]. In the hippocampus of IPF mice, only S1PR1 expression was significantly altered, with no significant changes detected in S1PR2–S1PR5. This selective upregulation may be explained by several factors. First, S1PR1 is uniquely sensitive to peripherally derived S1P. Under IPF condition, lung-derived S1P enters the circulation and likely first contacts and activates S1PR1 on blood–brain barrier endothelial cells [30], thereby initiating downstream signaling cascades. In contrast, S1PR2 and S1PR3 are predominantly localized to neuronal synapses and astrocytes and are relatively insensitive to fluctuations in peripheral S1P [54]. Second, previous studies have shown that, in a chronic unpredictable mild stress (CUMS) mouse model of depression, hippocampal neuronal S1PR3 expression is markedly reduced, and S1PR3 overexpression rescues depressive-like behaviors, indicating that S1PR3 acts in the opposite direction to S1PR1 [55]. In the IPF model, however, no such downregulation of S1PR3 was observed, suggesting fundamental differences in the regulation of the S1P receptor system between these two pathological states. Third, in an experimental autoimmune encephalomyelitis (EAE) model, the expression of S1PR1, S1PR3, and S1PR5 in the hippocampus is globally downregulated following EAE induction, reflecting complex dynamic changes in receptor expression [56]. Nevertheless, in the chronic low-grade IPF model, the selective activation of S1PR1 may reflect the specificity of receptor expression regulation in response to distinct pathological stimuli. Notably, S1PR1 is a receptor subtype widely expressed in neurons, astrocytes, oligodendrocytes, microglia and endothelial cells, and its signaling is coupled to Gαi proteins [35]. This broad distribution across multiple cell types positions S1PR1 as a key node through which peripheral signals regulate central nervous system function. S1PR1 is particularly sensitive to the upregulation of inflammatory mediators under neuroinflammatory conditions, and its aberrant activation can drive neuropathological processes by promoting microglial inflammatory responses and glial activation [36]. Altered S1PR1 expression has been associated with mood disorders, including depression and anxiety. In a rat model of depression, activation of the SP1/SK1/S1PR1 signaling pathway was closely linked to hippocampal neuroinflammation and impaired synaptic plasticity, and modulating S1PR1 levels ameliorated depressive-like behaviors [57]. Notably, the direction of S1PR1 dysregulation varies across pathological conditions. In the EAE model, EAE induction leads to global downregulation of S1PR1 and S1PR3 in the hippocampus, and FTY720 treatment results in complex dynamic changes in receptor expression without improving anxiety-like behaviors [56]. In contrast, in a chronic pain model, downregulation of S1PR1 in the hippocampal dentate gyrus is closely associated with memory impairment, and an S1PR1 agonist prevents cognitive deficits [58]. Furthermore, S1PR1 protein expression is significantly elevated in the dorsolateral prefrontal cortex of patients with schizophrenia compared with controls [59]. These findings indicate that the expression and functional effects of S1PR1 are highly disease- and brain region-specific. In the present study, we demonstrated that, in IPF mice, elevated serum S1P selectively activates hippocampal S1PR1, leading to depression- and anxiety-like behaviors, and that intervention with Fingolimod reversed these abnormalities and improved emotional behavior. This may be attributed to the widespread brain distribution and sensitivity to inflammation of S1PR1. These findings together constitute an important molecular basis for the remote regulation of hippocampal function by pulmonary fibrosis.

Upon ligand binding, S1PR1 regulates multiple downstream signaling pathways and molecules, including PI3K/Akt and MAPK/ERK signaling, Rho family GTPases, and phospholipase C (PLC) [30, 60–62]. The PI3K signaling pathway is one of the most critical signal transduction pathways in the central nervous system, is widely expressed in emotion-related brain regions, and participates in the regulation of various physiological processes such as neurogenesis, synaptic plasticity, and neuroinflammation. Numerous studies have confirmed that dysregulation of the PI3K/AKT pathway is closely associated with the pathogenesis of depression [63–65]. In the present study, we observed significantly elevated levels of p-PI3K in the hippocampus of IPF mice, indicating aberrant activation of this signaling pathway. This finding differs from the conventional view that the PI3K/AKT pathway is generally suppressed in depressive states. In fact, recent studies have suggested that the role of the PI3K/AKT signaling pathway in depression is complex, and its activation status may depend on the nature and duration of upstream stimuli as well as cell type-specific differences [66]. The elevated phosphorylation observed in our study may reflect a chronic stimulatory effect of persistently elevated S1P on the hippocampus via S1PR1 in IPF. PKA is a key serine/threonine protein kinase that regulates the function of various downstream effector molecules through phosphorylation. Aberrant activation of PKA is involved in regulating microglial inflammatory responses and cytokine production [67]. The PKA/CREB signaling pathway is also well recognized for its role in neuroplasticity and mood regulation. In various stress-induced models of mood disorders, the PKA/CREB signaling pathway is typically suppressed, a mechanism involving the regulation of BDNF and neuropeptide Y, as well as the modulation of GABA dysfunction, HPA axis dysregulation, and neuroinflammation [68, 69]. However, in the present study, we observed aberrant activation of the PKA/CREB pathway in IPF mice. This seemingly contradictory finding suggests that IPF-associated mood disorders may differ mechanistically from classical stress-induced depression. Moreover, the PI3K and PKA pathways in the hippocampus may act synergistically in IPF-induced mood disorders. These findings may reveal the mechanisms by which lung-derived S1P induces mood disorders and provide new insights into how this signaling pathway mediates hippocampal neuronal injury. Intervention with Fingolimod, a selective functional inhibitor of S1PR1, significantly suppressed activation of the above pathways and ameliorated emotional behaviors in IPF mice.

Ferroptosis is a recently identified form of regulated cell death characterized by iron-dependent accumulation of lipid peroxidation and inactivation of glutathione peroxidase 4 (GPX4) [70, 71]. In recent years, accumulating evidence has linked ferroptosis to the pathogenesis of depression. Dysregulation of the ferroptosis–mitochondria axis participates in depression through a vicious cycle of oxidative stress, energy deficit, and neuroinflammation [72], and targeting ferroptosis may represent an effective strategy for ameliorating depression [73–75]. These findings are highly consistent with the results of the present study. Although preliminary, we observed marked changes in the ferroptosis markers ACSL4 and GPX4 in the hippocampus of IPF mice. This form of cell death may serve as one of the pathological bases for hippocampal neuronal injury and mood disorders in IPF. Notably, our study also revealed a potential link between the PI3K/PKA/CREB signaling pathway and ferroptosis. The PI3K/Akt signaling pathway is closely involved in the regulation of ferroptosis [71], and the PKA signaling pathway can modulate ferroptosis through phosphorylation-mediated downstream effectors [76]. In the hippocampus of IPF mice, the PI3K/PKA/CREB pathway may drive emotional disorders by regulating hippocampal ferroptosis. More importantly, treatment with Fingolimod effectively reversed the changes in ACSL4 and GPX4 expression and alleviated anxiety- and depression-like behaviors in the IPF model, further supporting a critical role for ferroptosis in IPF-induced emotional disorders.

Fingolimod (FTY720) is an FDA-approved selective functional inhibitor of S1PR1 used for the treatment of multiple sclerosis, and its neuroprotective effects on the central nervous system have attracted considerable attention in recent years [77–80]. The present study is the first to apply Fingolimod to IPF-associated emotional disorders. Fingolimod treatment not only significantly ameliorated mood disorders but also inhibited glial activation and neuroinflammation and reduced cell death in the hippocampus of IPF mice. These findings are highly consistent with previous reports on the antidepressant effects of Fingolimod [81, 82]. This observation has important clinical translational implications, suggesting that Fingolimod or its analogs may serve as candidate drugs for treating emotional disorders in IPF patients. However, it is worth noting that the effects of Fingolimod on the central nervous system are not uniform across all contexts. For instance, one study reported that although Fingolimod effectively reduced hippocampal inflammation in an experimental autoimmune encephalomyelitis (EAE) model, it failed to improve anxiety-like behaviors [56]. The impact of Fingolimod on emotional behavior may depend on factors such as disease type and dosing regimen.

The concept of the “lung–brain axis” has gained increasing attention in recent years. Accumulating evidence suggests that lung–brain interactions play a role in the development and progression of diseases and comorbidities affecting both organs [83, 84]. The neuroinflammation and hypoxia-induced injury have been identified as key factors that mediate this axis [85–88]. Airborne particulate matter can activate inflammatory responses in alveolar epithelial cells and subsequently induce microglial activation in the brain [89]. Chronic obstructive pulmonary disease (COPD), asthma, obstructive sleep apnea (OSA), and pulmonary infections may contribute to cognitive decline through systemic inflammation, immune crosstalk, hypoxic injury, and air-pollutant-induced neurotoxicity [90]. Most of these studies on pulmonary diseases and brain dysfunction have focused on cognitive impairment and its underlying mechanisms. Numerous reports have also described mood disorders associated with pulmonary pathologies, including IPF [21, 91, 92]. However, the specific molecular mechanisms linking chronic fibrotic lung disease to mood disorders via the lung–brain axis have not been systematically investigated. In this study, via the multi-omics approach, we discovered for the first time that a single metabolite, S1P, derived from IPF lung tissue, induces depression. Our findings provide novel and concrete evidence for the molecular mechanisms of the lung-brain axis. The signaling pathway model we propose “fibrotic lung tissue releases S1P → transport via the bloodstream → activation of hippocampal S1PR1 → phosphorylation of the PI3K/PKA/CREB pathway → neuroinflammation and ferroptosis → altered emotional behaviors” delineates a complete causal chain from a peripheral organ to the central nervous system and from molecular signals to behavioral phenotypes. This model provides a paradigm and reference for other neurological disorders associated with pulmonary conditions.

Several limitations of the present study should be acknowledged. First, the study was primarily conducted using a mouse model of IPF induced by bleomycin. Although this is one of the most commonly used animal models in IPF research, it does not fully recapitulate all pathological features of human IPF, such as progressive irreversible fibrosis and heterogeneous lesion distribution. Therefore, our findings need to be validated in additional IPF animal models and translated to IPF patients to assess the clinical feasibility of the use of S1P and related signaling molecules as biomarkers and therapeutic targets. Second, the hypothesis that elevated Mfsd2a expression in the lungs of IPF patients leads to increased S1P efflux from the lungs requires further experimental confirmation. Third, the use of pharmacological approaches in this study has inherent limitations. We administered SKI-V intraperitoneally to inhibit Sphk1 function in the lungs. Although this approach successfully reduced serum S1P levels, it cannot completely exclude the systemic effects of SKI-V on other tissues and organs. Future studies using genetic approaches, such as conditional knockout of Sphk1 in lung tissue or specific knockout of S1PR1 in the hippocampus, could further elucidate the role of the Sphk1/S1P/S1PR1 axis in IPF-induced mood disorders. A similar issue applies to the use of Fingolimod (FTY720). Fingolimod is an immunomodulatory agent, and its effects on the systemic immune system may indirectly contribute to the regulation of emotional behavior. Although the present study focused on local signaling pathway changes in the hippocampus, the possible contribution of altered peripheral immune cell function cannot be completely ruled out. Future studies employing central local administration (e.g., intracerebroventricular injection or local hippocampal microinjection) could help distinguish the direct central effects of Fingolimod from its peripheral indirect effects. Nevertheless, our pharmacological results provide preliminary validation of our conclusions. Fourth, the role of the PI3K/PKA/CREB signaling pathway and its causal relationship with ferroptosis were not further validated by gain- or loss-of-function experiments, nor did we identify the specific cell types in which hippocampal S1PR1 upregulation and signaling pathway changes occur. Future studies using immunofluorescence staining, single-cell sequencing, and in vitro experiments are warranted. Finally, the observation endpoints of this study were focused primarily on short- to medium-term (weeks) effects, and the long-term efficacy and safety of the interventions were not evaluated. Given that IPF is a chronic progressive disease requiring long-term management, the safety and efficacy of pharmacological interventions need to be assessed in further studies. Nonetheless, our study provides substantial evidence that fibrotic lung-derived S1P is indeed involved in the development of emotional disorders.

## Conclusion

Our study demonstrates that fibrotic lung-derived S1P enters the bloodstream. This supraphysiological level of circulating S1P activates hippocampal S1PR1, which triggers the PI3K/PKA/CREB signaling pathway, leading to microglial and astrocytic activation, synaptic dysfunction, neuroinflammation, and ferroptosis, thereby driving emotional disorders. Pharmacological inhibition of Sphk1, the key enzyme in pulmonary S1P synthesis, reduces serum S1P levels and alleviates emotional disorders in IPF mice. Similarly, selective inhibition of hippocampal S1PR1 with Fingolimod ameliorates neuroinflammation and ferroptosis, thereby relieving emotional disorders in IPF mice (Figure 9). These findings demonstrate that a metabolite derived from fibrotic lung tissue can induce emotional disorders, providing new potential intervention targets for emotional disorders in IPF patients. Moreover, this study offers novel perspectives and a reference for research on other pulmonary disease-related neurological diseases.

**Fig. 9.**
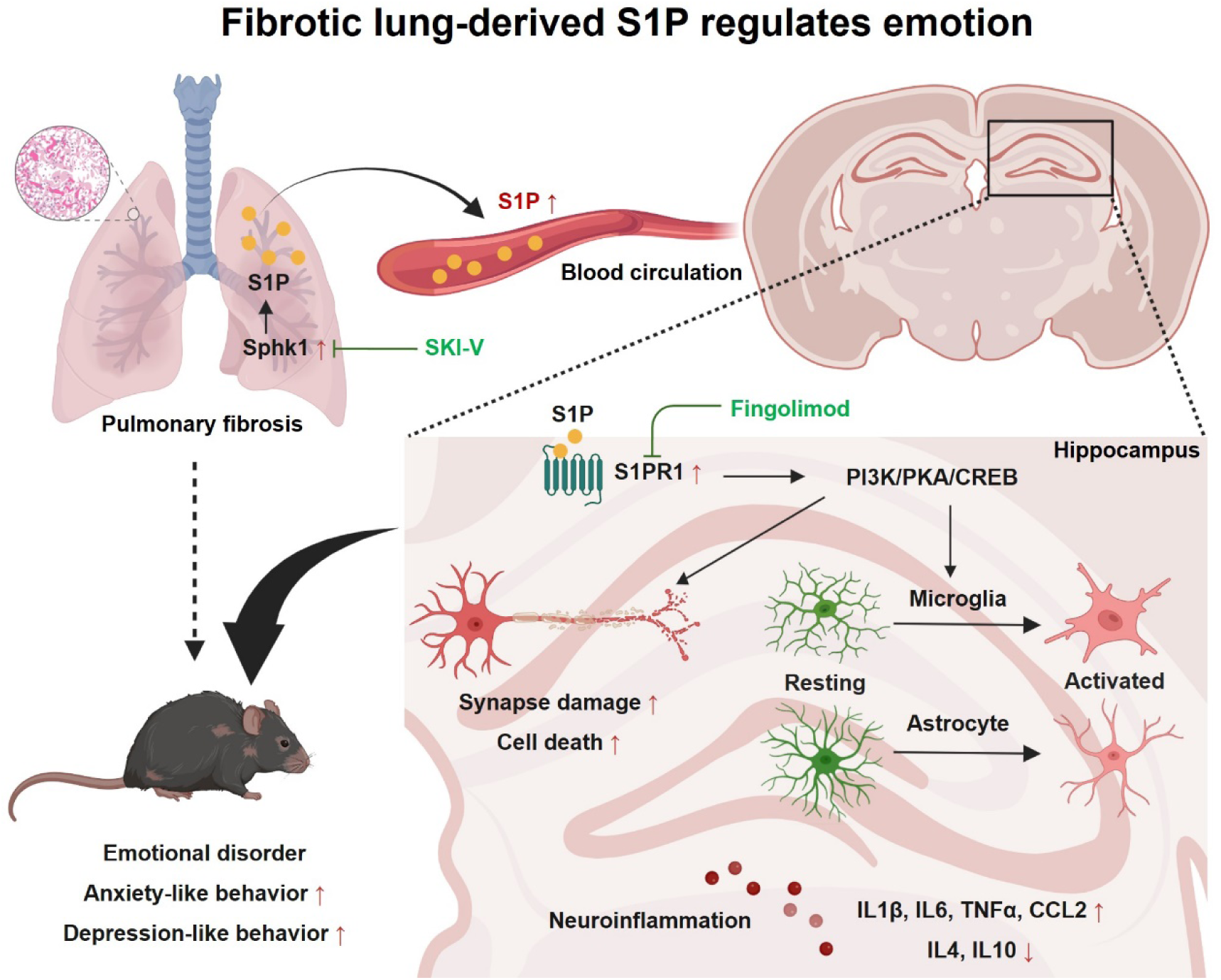
S1P derived from fibrotic lungs drives emotional disorders and the underlying mechanism. S1P originating from the fibrotic lungs enters the bloodstream. This supraphysiological level of circulating S1P binds to hippocampal S1PR1, which triggers the PI3K/PKA/CREB signaling pathway, leading to microglial and astrocytic activation, synaptic dysfunction, neuroinflammation, and ferroptosis, thereby driving the onset of mood disorders. Pharmacological inhibition of pulmonary Sphk1 reduces serum S1P levels and alleviates mood disorders in IPF mice. Similarly, functional inhibition of hippocampal S1PR1 with Fingolimod ameliorates neuroinflammation and ferroptosis, thereby alleviating emotion disorders in IPF mice. The graphical abstract was made in Biorender (https://app.biorender.com/).

## Supplementary Information

Supplementary Material 1: Fig.S1 The S1P content in the hippocampus of pulmonary fibrosis mice remained unchanged, related to Fig.7.

Supplementary Material 2: Excel file listing antibodies and qPCR primers.

Supplementary Material 3: uncropped Western blotting bands.

## Acknowledgements

The authors thank the Applied Neuroscience Research Center, School of Life Science and Technology, Henan Medical University, for providing the behavioral experimental platform.

## Author contributions

Z.X. and YL.L. conceived the idea. Z.X. designed the experiments, provided funding and wrote the manuscript. D.X. and K.L. conducted the experiments and processed the data. Y.Y. and X.H. analyzed the data. F.Y. provided some materials. W.N. provided funding. J. D., YH.L. and P.Z. supervised, proofread, and edited the manuscript. R.G., YL. L. and L.J. supervised and provided funding. All the authors have reviewed the manuscript.

## Data availability

The original RNA-seq data have been uploaded to the NCBI database. The original metabolome data have been uploaded to the Metabolomics Workbench. These data will be made publicly available one year after publication of the article. The other data used and analyzed in the present study are available from the first and corresponding authors on reasonable request.

## Ethics approval and consent to participate

All animals were used in accordance with the Code of Ethics of the World Medical Association. All experimental protocols and procedures were approved by the Ethics Committee of Henan Medical University.

